# Type II topoisomerase substrate geometry revealed through combined experiment and computation

**DOI:** 10.64898/2026.09.04.749274

**Authors:** Mihirkumar N. Prajapati, Yeonee Seol, Jonathan Silver, Parth R. Desai, Siddhartha Das, Keir C. Neuman

**Author notes:** Corresponding author: Dr. Keir C. Neuman **Email:**. **Author Contributions:** M.N.P., S.D., and K.C.N. conceptualized the study. M.N.P. wrote the manuscript, and all other authors edited it. Y.S. performed the experiments. M.N.P. and P.R.D. designed the LAMMPS simulation, and M.N.P. performed the simulations. M.N.P. and J.S. analyzed the simulation and experimental results. **Competing Interest Statement:** None declared.

## Abstract

Type II topoisomerases (topo IIs) are essential enzymes that regulate DNA topology through a strand-passage mechanism in which a duplex DNA (transfer-segment) is passed through a transiently cleaved second duplex DNA (gate-segment). Biochemical and structural approaches have revealed critical details of the binding and cleavage of the gate-segment DNA. However, capture of the transfer-segment DNA has proven more difficult to resolve due to the transient nature of the interaction. Nonetheless, selection of the transfer segment with a specific conformation or orientation relative to the gate segment is predicted to govern aspects of topo II activity, including chiral discrimination and the ability to reduce topological complexity below equilibrium. To determine the conformation and orientation of the transfer-segment relative to the gate segment, we combined experimental single-molecule measurements of topo II unlinking a single DNA crossing with Brownian dynamics simulations of the DNA crossing. By correlating the unlinking rate with the geometric features of the DNA crossing, we obtain the complete three-dimensional preferred crossing geometry for strand passage. Strikingly, the preferred crossing geometries for *Escherichia coli* topoisomerase IV and *Methanosarcina mazei* topoisomerase VI are distinct and provide structural models of the DNA synapse selected for strand passage along with a mechanistic basis for their differing activities and biological functions. The approach we develop is generalizable, providing unique insights into the kinetic selection of DNA synapse structure.

**Significance Statement:** Type II topoisomerases resolve DNA entanglements by passing one DNA duplex through a transient break in another, but how these enzymes select the three-dimensional geometry of the DNA segments they act on has remained unresolved because the interaction is too transient to capture structurally. We developed an approach combining single-molecule measurements with Brownian dynamics simulations to determine the crossing geometry each enzyme kinetically selects. Applied to two distinct topoisomerases, this approach revealed different preferred geometries that explain long-standing differences in how these enzymes sense DNA handedness and challenge a leading model for how topoisomerases simplify DNA tangles beyond what thermodynamics alone would predict. This kinetics-based strategy is broadly applicable to other enzymes that resolve DNA or RNA synapses.

## Introduction

Type II topoisomerases (topo IIs) are essential enzymes that regulate DNA supercoiling and resolve knots and catenanes to maintain the integrity and topological homeostasis of the genome (1–3). They function by transiently cleaving both strands of one DNA segment, the gate-segment (G-segment), and passing a second segment, the intact transfer-segment (T-segment), through the break (4). Although the double-strand break is highly vulnerable (5–9), topo IIs govern the reaction such that cleavage occurs under specific conditions and is followed by rapid religation (10, 11). This ATP-dependent strand-passage activity allows topo IIs to alter DNA topology; supercoiling and knotting via intra-segmental transfer or decatenation and catenation via inter-segmental transfer (11–13). These activities are essential during DNA metabolic processes, including DNA replication, transcription, and chromosome segregation, underscoring the importance of topo IIs across all domains of life (11–13). Type II topoisomerases comprise two subclasses—type IIA and type IIB—that share mechanistic similarities but exhibit notable structural, biochemical, and evolutionary differences (4, 14).

Type IIA topoisomerases display remarkable ability to operate beyond thermodynamic equilibrium (15), actively reducing DNA topological complexity, which is not observed in type IIB topoisomerases (16, 17). Rybenkov et al. (15) showed that type IIA topoisomerases (except gyrase, a type IIA topoisomerase) simplify DNA topology to yield steady-state concentrations of unlinked and unknotted molecules below thermodynamic equilibrium, and distributions of relaxed circular DNA topoisomers with variances below thermodynamic equilibrium. Since topo II strand passage consumes energy in the form of ATP hydrolysis, below-equilibrium simplification does not violate the second law of thermodynamics (15). However, it raises fundamental mechanistic questions: how do these enzymes sense global DNA topology while acting at the local scale.

Structural and biochemical studies revealed the importance of G-segment bending in regulating cleavage and have provided insights into the structural features that accommodate T-segment passage (18–23). The binding of the topoisomerase to the G-segment is relatively stable, mediated by extensive protein-DNA contacts, including the covalent attachment between the topo II catalytic tyrosines and the 5’ cleaved ends of the G-segment (22, 23). However, interactions between T-segment and topoisomerase are transient, though recent structural studies have identified a few weak interactions between the T-segment and gyrase (24, 25). The consensus view is that after the binding of the topoisomerase to the G-segment, ATP binding closes the topo II N-gate, capturing the T-segment and passing it through the cleaved G-segment into the C-gate, which is subsequently opened to release of the T-segment in conjunction with the later steps in the ATPase cycle (1, 13, 26, 27).

Despite the transient interactions of the T-segment with the topo II, the process of T-segment capture, and particularly the conformational and/or geometric selection of the T-segment, is postulated to underlie specific activities of topo IIs (28–36). Chiral discrimination, the preferential relaxation of positively versus negatively supercoiled DNA, has been attributed to selection of a particular G- and T-segment juxtaposition geometry (28, 29). *Escherichia coli* Topoisomerase IV (*E. coli* Topo IV), a type IIA topo, preferentially relaxes positive writhe, due primarily, but not entirely, to dramatic processivity differences (highly processive on positive supercoils, while perfectly distributive on negative supercoils), (21, 28). On the other hand, *Methanosarcina mazei* topoisomerase VI (*M. mazei* Topo VI) a type IIB topo, efficiently unlinks catenanes and braids while slowly and distributively relaxing supercoils suggesting that it is an intrinsically preferential decatenase (29, 37). Topo IV relaxes positive supercoils at least 20-fold faster than negative supercoils (28). In contrast, Topo VI relaxes positive supercoils a modest ~2-fold faster than negative supercoils (29). Whereas the average preferred crossing angle for both enzymes was found to be similar (just below 90°) (28, 29), topo IV and topo VI showed large differences in enzymatic rates and processivity dependent on the DNA topological substrates suggesting the simple single-crossing angle description of the DNA crossing geometry does not fully capture the detailed configurations of the crossing synapses and the mechanistic details of geometric T-segment selection.

Selection of a particular configuration of G-and T-segments has also been proposed to explain the below-equilibrium topology simplification activity of type IIA topos (30–36). Modeling studies suggest that topology simplification below thermodynamic equilibrium levels can be achieved if topo II selectively unlinks G- and T-segments sharply bent towards each other in a “hooked juxtaposition” (Fig. 1A) (30–36). A “fully-hooked” juxtaposition, with bent G- and T-segments, produces greater topology simplification than a “half-hooked” juxtaposition (30–36). Both the chiral discrimination model (28, 29) and the hooked juxtaposition (HJP) model (30–36) are predicated on topo II selectively acting on a particular G- and T-segment geometry established before topo II binding. However, the extent to which topo IIs preferentially capture T-segments with a particular conformation, and the full three-dimensional geometry of the preferred juxtaposition, remain unresolved.

**Figure 1.**
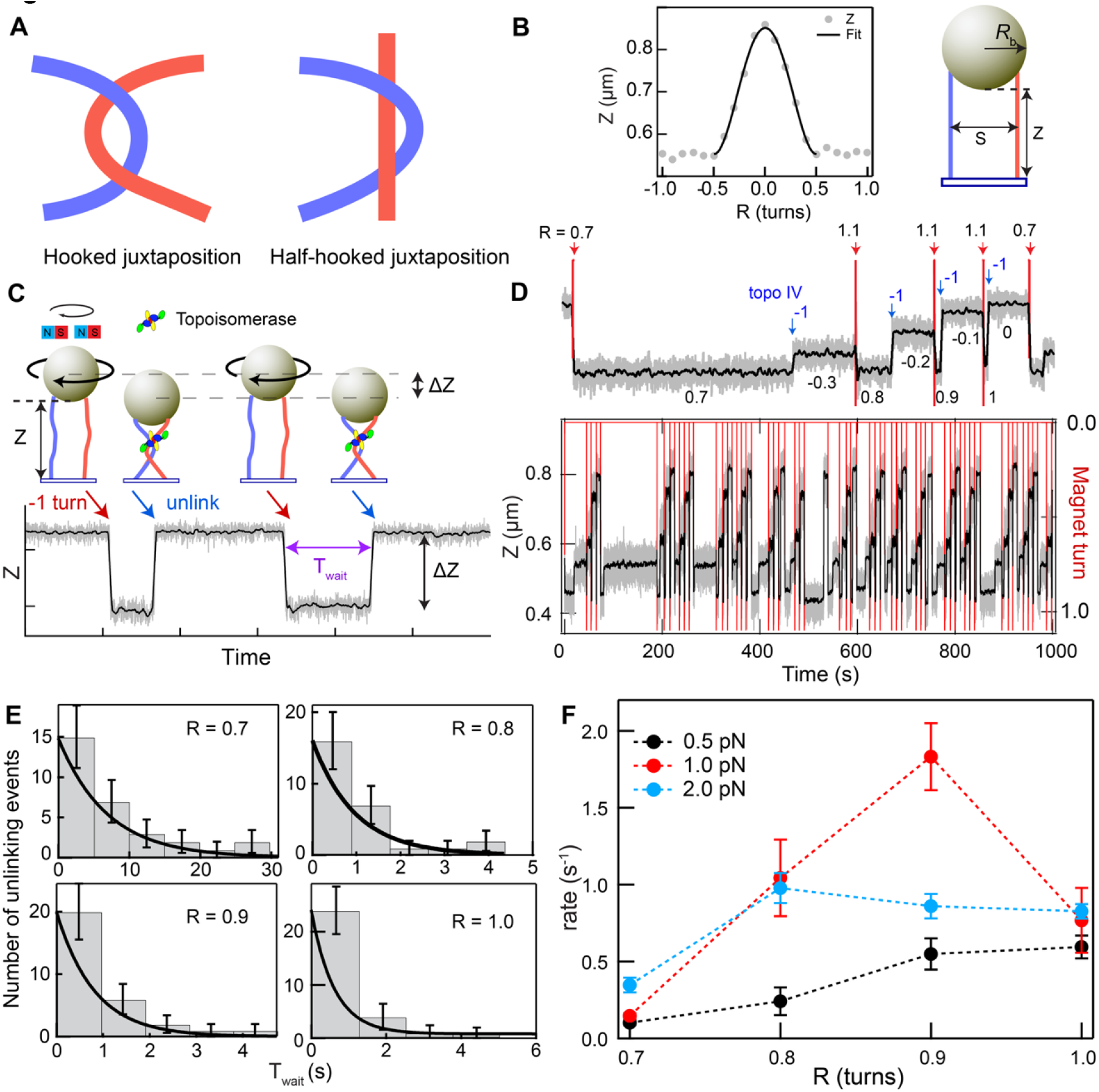
Single-molecule Topoisomerase IV unlinking assay. **A.**Schematics of hooked and half-hooked juxtapositions. **B**. Characterization of DNA tether geometry. The double tether comprises two 3.6 kb DNA molecules attached between the surface and a streptavidin-coated magnetic bead (grey). To obtain the tether geometry, the bead height, Z, was measured as a function of the bead rotation from −0.5 to 0.5 rotations in increments of 0.1 turns (gray filled circles). The bead height over the range of −0.5 to 0.5 rotations was fitted with Eq. 2 (black solid line, methods) to obtain the spacing between the DNA molecules, S, and the bead height at zero rotation, Z_0_. **C**. Cartoon of experiment (not to scale). 1 µm magnetic bead (beige) is tethered to the surface by two 3.6 kb torsionally unconstrained dsDNA molecules (blue and red). A link between the DNA molecules is generated by rotating the bead (black arrow) using a magnet assembly above, which decreases the bead height by ΔZ. Strand passage mediated by a topo IV unlinks the DNA (blue arrow), leading to an increase in bead height, Z. After a brief system equilibrium time, the process is automatically repeated. Extension of the double tether, Z-trace, shows the decrease in extension associated with introducing the crossing and the subsequent increase in extension when the unlinking activity of topo IV resolves the crossing. The time between these events is defined as the waiting time (T_wait_). DNA extension was measured by video tracking the bead in real-time at 200 Hz. **D**. Example trace showing topo IV unlinking a series of DNA crossings ranging from 0.7 to 1 imposed turns. Inset: Topo IV unlinking at different imposed turns (0.7, 0.8, 0.9, and 1). Except for the 0.7 turn, 1.1 turns were added to impose 0.8 to 1 turn as topo IV unlinking changes the turn number by −1 (blue arrows). For example, for the 0.7 turn condition, a single unlinking event leaves a residual turn of −0.3. So, in order to introduce 0.8 turn in the subsequent step, 1.1 turn should be added. The numbers under the trace show the number of turns imposed on the DNA. **E**. Representative T_wait_ distributions for each imposed rotation (R) (S = 720 nm; F = 1 pN) with exponential fits to obtain unlinking rate for each rotation. **F**. Topoisomerase IV unlinking rate as a function of bead rotation, R, for three different forces with a DNA spacing, S, of 720 nm (F=0.5 pN: Black; 1 pN: Red; 2 pN: Blue). The error bars correspond to the standard error of mean.

To directly test these models and address the gaps in our mechanistic understanding of geometric selection of T-segment, we require an approach that can determine the geometry of the G- and T-segments acted on by topo II prior to strand passage, and indeed prior to enzyme binding. Although the structure-based approaches provide detailed T-segment-bound topo II structures for DNA gyrase (24, 25), they might reveal how segments are reconfigured on binding rather than the geometry of the G- and T-segment juxtaposition selected for binding, which is potentially more important for understanding how topo IIs select juxtaposed DNA segments for strand passage.

To obtain the geometry of the G- and T-segment synapse selected by topo IIs for strand passage, we employ Brownian dynamics simulations that mimic the experimental configurations from the current and a previous experimental study (29) and investigate the relationship between strand-passage activity and crossing geometry. We create a single DNA crossing *in vitro* using a magnetic tweezers assay in which two DNA molecules are attached to a magnetic bead and vary the crossing geometry by rotating the bead to different extents at different applied forces (38). Strand-passage activity is quantified as the unlinking rate of each crossing geometry. Since simulations allow detailed characterization of the G- and T-segments synapse (crossing geometry) beyond the single “crossing angle” obtained in previous studies (28, 29), we calculate an expanded set of geometric parameters that fully define the three-dimensional geometry of the synapse from simulations. By correlating the simultaneous, or joint, probability of obtaining particular geometric parameters with the experimentally measured enzyme unlinking rates, we identify the preferred crossing geometry defined by the geometric parameters that maximize the correlation.

Using this approach, we demonstrate that *E. coli* topo IV and *M. Mazei* topo VI have similar but distinct preferences for juxtaposed G- and T-segment geometries, consistent with their respective chiral discrimination and processivity behaviors. Additionally, our results revealed that the preferred geometry of topo IV more closely resembles positive writhe with gently bent G- and T-segments, inconsistent with the sharp symmetric bending predicted by the HJP model to produce the degree of topological simplification achieved by topo IV. More generally, we demonstrate a robust approach to obtain the geometry of a kinetically selected DNA synapse that is applicable to synapse-resolving enzymes.

## Results

### Topo IV unlinking rate exhibits a non-monotonic dependence on imposed crossing geometry

We employed a single-molecule DNA unlinking assay to measure the *E. coli* topo IV unlinking rate as a function of the geometry of the juxtaposed DNA segments in a single crossing (28, 38–40). A single DNA crossing was generated by rotating a magnetic bead tethered to the surface by two DNA molecules (Fig. 1C). The height change, calculated from purely geometric considerations, depends on the length of the DNA (L), the distance between the two DNA molecules (S), the force on the bead (F), and the imposed bead rotation (R) (methods). We determined the crossing geometry and ensured that the DNA molecules were parallel by fitting the DNA extension, Z, as a function of bead turns with Eq. 2 as described in the methods section (Fig. 1B).

Experiments consisted of imposing a single crossing (0.7 to 1 turn) between the two DNA molecules, producing a decrease in bead extension. In the presence of topo IV and ATP, imposed crossings were unlinked after a variable waiting time T_wait_, resulting in an increase in the bead height (Fig. 1C). A crossing was re-imposed after each unlinking event, and the process was repeated for 30 or more unlinking events for each imposed rotation. Bead height traces (Fig. 1D) were analyzed using a custom step-finding algorithm to identify the waiting time, T_wait_. T_wait_ values were binned and the histogram was fitted with a single exponential to obtain the unlinking rate for each imposed rotation. Fig. 1E shows representative T_wait_ distributions and corresponding exponential fits of the unlinking rates for different values of bead rotation (R = 0.7 to 1 turns) in the presence of 2 nM topo IV, above the dissociation constant, K_d_ of ~0.2 nM, measured under similar conditions (41) and saturating (1 mM) ATP. The unlinking rate generally, but not uniformly, increases with increasing rotation for all three applied forces (F = 0.5, 1, and 2 pN) (Fig. 1F).

Experiments were performed with 12 double tethers spanning a range of segment spacings (S = 440 nm to 960 nm), each measured at a range of forces (0.5, 1, and 2 pN), and bead rotations of 0.7 to 1 turn in 0.1 turn increments imposing positive writhe. At F = 0.5 pN (Fig. S1 top panel), unlinking rates are generally lower for all segment spacings than at higher forces. At F =1 pN (Fig. S1 middle panel), unlinking rates exhibit a non-monotonic dependence on rotation for most segment spacings, suggesting an optimal crossing geometry for topo IV unlinking. At F = 2 pN (Fig. S1 bottom panel), unlinking rates are generally lower than at 1 pN. Notably, the non-monotonic variation of unlinking rate with bead rotation at all three forces contrasts with the monotonic increase observed previously with longer (5 kb) DNA molecules (38).

These results reveal an intricate interplay between segment spacing, applied force, and bead rotation in governing topo IV unlinking rate. However, the fluctuating microscopic geometric features of the DNA crossings that underlie this behavior cannot be directly obtained from the experimental data.

### Brownian dynamics simulations reveal crossing geometry distributions

Single-molecule experiments quantify enzyme activity under different crossing conditions, defined by different combinations of spacing (S) between the two DNA molecules, the applied force (F), and the imposed number of turns (R). Increasing S, F, and R each increase the degree of “hookedness”, the extent of bending of the two juxtaposed segments towards each other (Fig. 1A and 2A-C), but the three dimensional characterization of each crossing configuration and their fluctuations due to Brownian motion could not be calculated from the experimental crossing conditions. To quantify the crossing geometry and the distributions of geometric parameters due to Brownian motion, we performed Brownian dynamics simulations of each DNA double tether with the experimental values of DNA length and separation, force, and rotation (Methods).

The two DNA in the simulations were configured to reproduce the experimental double-tether geometry. Two linear charged polymers were arranged in a parallel configuration at a fixed separation. One end of the double-molecule was rotated about its long axis while the other end was kept fixed. Representative simulation snapshots (Fig. 2 A-C) illustrate three of the 83 simulated geometries, spanning low (I), medium (II), and high (III) levels of hookedness. At low hookedness (smaller values of S, F, R) the crossing is loose and poorly localized whereas at high hookedness (larger values of S, F, R) the crossing is tight and localized (Movie S1).

**Figure 2.**
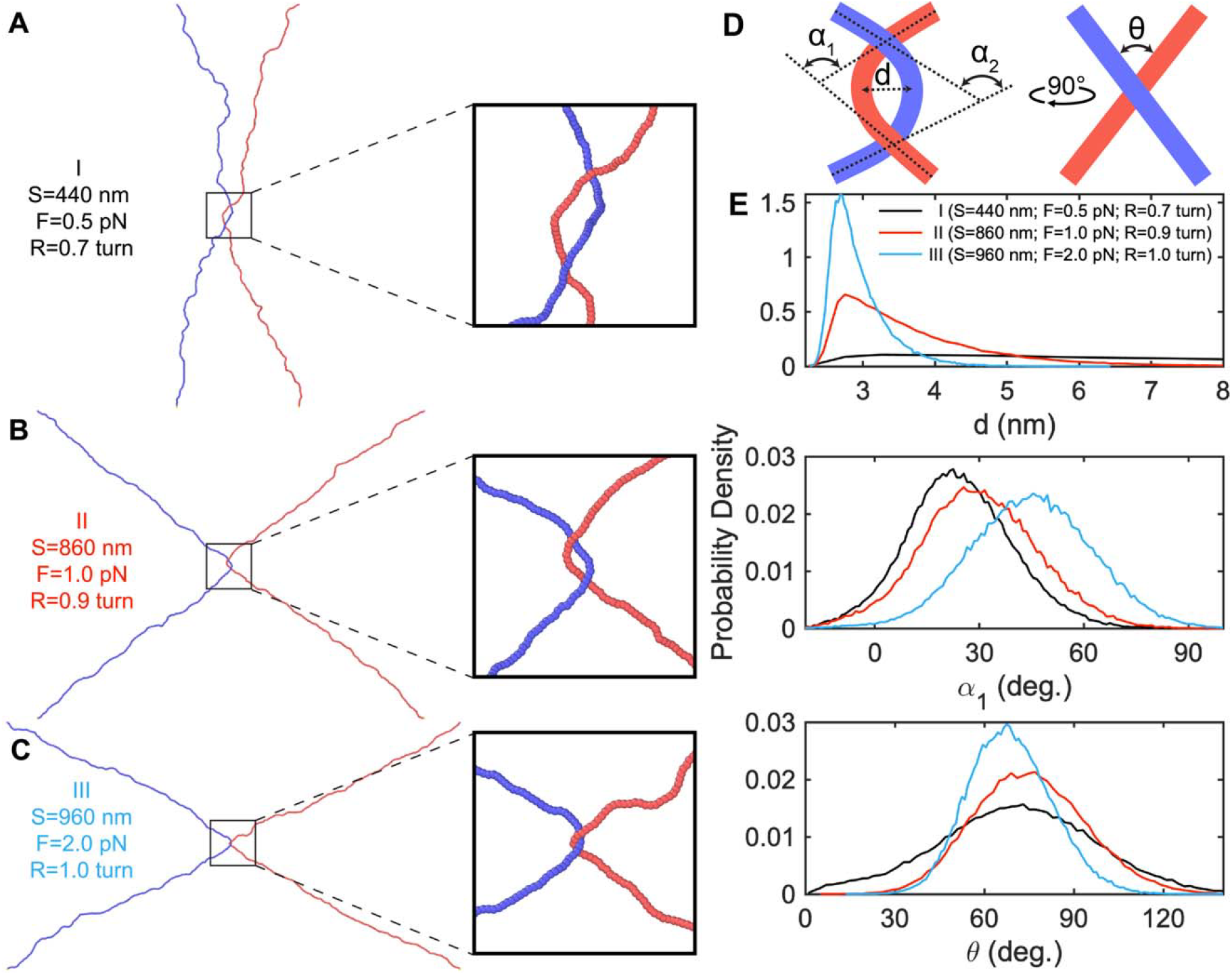
**A-C**. Representative snapshots of simulated crossings illustrating different degrees of hookedness. The left column shows simulation snapshots, and the right column shows expanded views centered on the crossing. **A**. Low hookedness crossing: S = 440 nm, F = 0.5 pN, R = 0.7 turn (252°). **B**. Medium hookedness crossing: S = 860 nm, F = 1 pN, R = 0.9 turn (324°). **C**. High hookedness crossing: S = 960 nm, F = 2 pN, R = 1 turn (360°). The thermal motions of crossings at these representative cases can be seen in Movie S1. **D**. Schematic illustrating crossing geometry parameters. Juxtaposition distance (*d*) is defined as the distance at the closest approach. The bend angles (*α*_1_, *α*_2_) are defined as the angle between the terminal tangent vectors for a given segment centered on the point of closest approach. The crossing angle (*θ*) is defined as the angle between the planes defined by the two segments. Right schematic is the side view of the schematic on the Left. The detailed calculations and illustrations of crossing geometry parameters are found in methods and Fig. S2. **E**. Probability density distributions of crossing parameters *d* (juxtaposition distance), *α*_1_ (bend angle), and *θ* (crossing angle) at three different levels of hookedness. The distributions are color-coded for each tuple ({S, F, R} = {440 nm, 0.5 pN, 0.7 turn}: Black; {860 nm, 1 pN, 0.9 turn}: Red; {960 nm, 2 pN, 1 turn}: Blue).

### Crossing geometric parameters exhibit monotonic trends with increasing rotation

At each simulation timestep, the crossing geometry is characterized by four parameters: juxtaposition distance (d), bend angles of each segment (α_1_ and α_2_), and crossing angle (θ) (Fig. 2D, Fig. S2, methods). Probability density distributions of the crossing parameters for the three example configurations are shown in Fig. 2E. The distributions of the crossing geometry parameters shift systematically with hookedness level (Fig. 2E). The *d* distributions become narrower and shift to smaller values with increasing values of S, F, and R, reflecting that more compact and stable crossings are expected as segment spacing, tension, and degree of wrapping increase. The α_1_ and α_2_ distributions shift to larger values with increasing levels of hookedness, consistent with increased segment bending. For most analysis, α_1_ serves as the representative bend angle as α_1_ and α_2_ have similar distributions and identical mean values (Fig. S3A). However, they are treated as separate parameters for subsequent data analysis as their values at individual time steps are uncorrelated (Fig. S3B). The θ distributions narrow but their means do not change significantly with increasing values of S, F, and R.

Figs. S4–S6 show mean crossing geometry parameters as a function of bead rotation for different strand separation distances at F = 0.5, 1 pN and 2 pN, respectively. All three mean parameters, <d>, <α_1_> and <θ>, vary monotonically with bead rotation.

### Individual crossing parameters are poorly correlated with unlinking rate

Under the assumption that the topo IV unlinking rate depends on the DNA crossing geometry, we first asked if any individual mean crossing parameter predicts unlinking rate. However, none of the mean crossing parameters were well-correlated with topo IV activity. The Pearson correlation coefficient between topo IV unlinking rate and <*d*>, α_1_, and (θ) were −0.37, 0.03, and −0.51, respectively (Fig. S7). The monotonic trends of the mean geometric parameters (Fig. S4–S6) thus cannot account for the non-monotonic unlinking rates, indicating that topo IV activity is not governed by the average value of any single geometric parameter.

### Joint-probability analysis to obtain the preferred crossing geometry

The non-monotonic topo IV activity (Fig. S1) when compared with the monotonic trends of mean crossing parameters as a function of bead rotation (Fig. S4–S6) suggests that topo IV preferentially acts on crossings with a specific geometry defined by multiple crossing parameters. In this scenario, the unlinking rate would be proportional to the probability of the parameters associated with the preferred crossing geometry occurring simultaneously. To test this model, we consider the probability of the simultaneous occurrence of the crossing geometry parameters (*d*, α_1_,α_2_, and θ) obtained from simulations and calculate the joint probability:

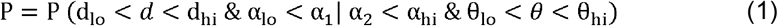

where P is the fraction of simulation timesteps for which all the conditions are simultaneously satisfied for a given configuration of S, F, R. The AND condition is represented by “&” and the OR condition is represented by “|”. “lo” and “hi” suffixes denote the lower and upper bounds, respectively, of a given crossing parameter. The OR condition on α_1_ or α_2_ removes the symmetry constraint on the crossing by allowing either α_1_ or α_2_ to satisfy the bend angle condition, which is consistent with potentially different interactions of the enzyme with the G- and T-segments at the crossing.

We then correlate this simultaneous probability calculated for each configuration with the associated unlinking rate. This approach allows us to identify the preferred crossing geometry by systematically varying the values of the selected geometric parameters to maximize the correlation between the simultaneous probability of observing the selected parameters in the simulations and the measured unlinking rate. In other words, in the multidimensional space of crossing parameters, we look for a region that is most correlated with enzyme activity.

We calculated correlation coefficients between enzyme activity and joint probabilities for 2.4×10^6^ different parameter combinations (d_lo_, d_hi_, α_lo_, α_hi_, θ_lo_, θ_hi_). The maximum correlation between the topo IV unlinking rate and the joint probability was 0.74 for the parameters: 3.6 nm < *d* < 3.9 nm, 35° < α_1_ | α_2_ < 39°, 55° < < 56°. Fig. 3A illustrates this optimal region in the (*d*, α, θ) parameter space. Fig. 3B shows a rendering of the corresponding preferred crossing geometry for topo IV.

**Figure 3.**
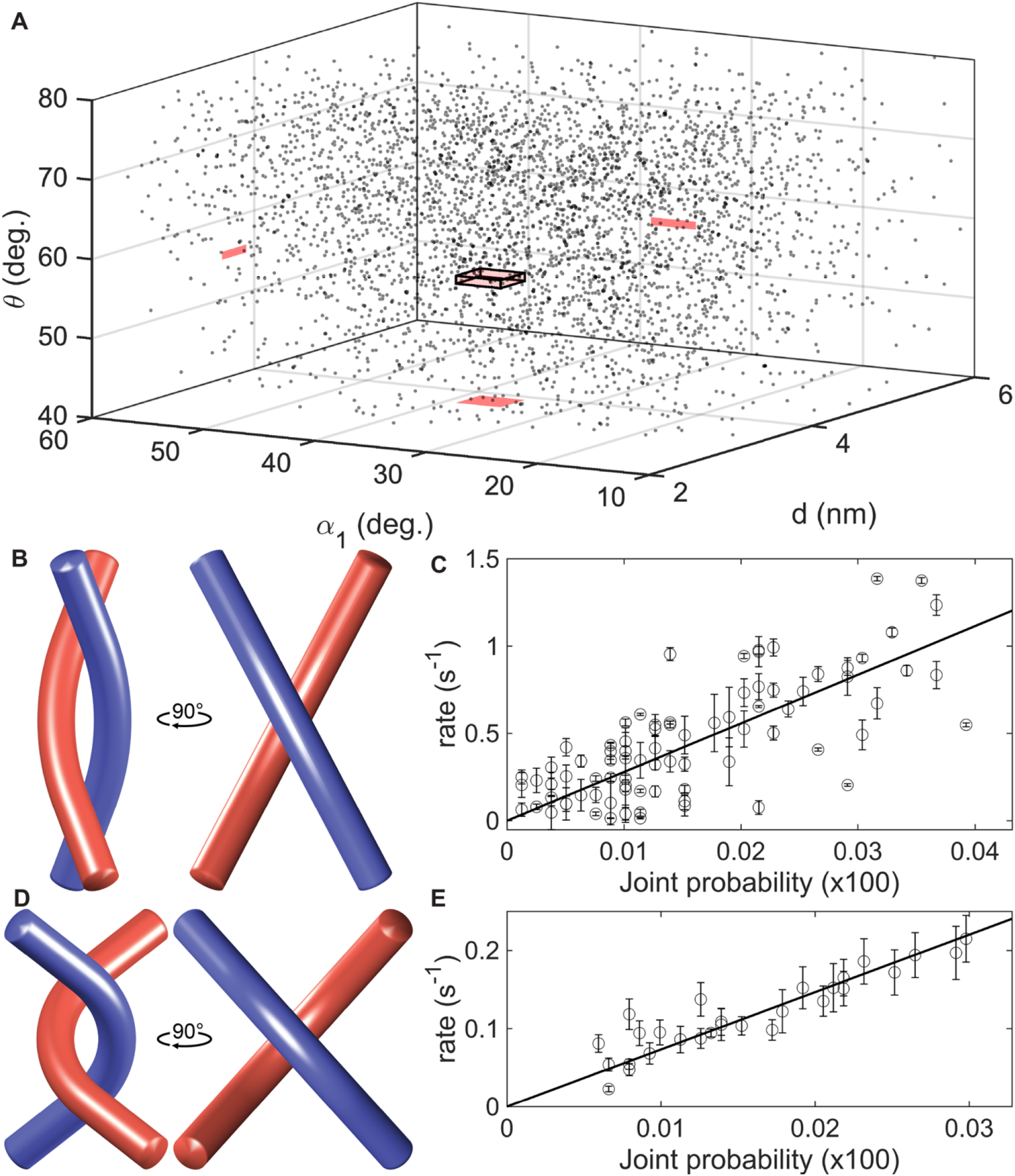
**A**. Representation of the joint probability volume in *d, α, θ*. The gray dots represent *d, α,θ*, for every 100^th^ timestep from a simulation for a given configuration. The optimum correlation (0.74) with topo IV unlinking rate was obtained for the range of parameters defined by the red box (3.6 nm < *d* < 3.9 nm & 35° < *α*_1_ | *α*_2_ < 39° & 55° < *θ* < 56°). The red rectangles represent the projection of the box on the respective planes. The joint probability is calculated as the number of points within the volume of the box divided by the total number of simulation points. **B**. Preferred DNA crossing geometry for topo IV: *d* = 3.75 nm, *α*_1_ = 37°, *α*_2_ = 37°, *θ* = 55.5°. **C**. Topo IV unlinking rate plotted as a function of the joint probability obtained with the maximum correlation geometric parameters. The line shows a linear fit (with 0 intercept, slope = 27.9 ± 1.4 S^-1^, and reduced chi-square, 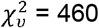). **D**. Preferred DNA crossing geometry for topo VI: *d* = 4.1 nm, *α*_1_ = 87°,*α*_1_ = 87°, *θ* = 83°. **E**. Topo VI unlinking rate plotted as a function of the joint probability obtained with the maximum correlation geometric parameters. The line shows a linear fit (with 0 intercept, slope = 9.4 ± 0.3 S^-1^, and reduced chi-square, 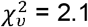). Error bars correspond to the standard error of the mean.

A scatter plot of unlinking rate as a function of the preferred crossing geometry joint probability and a linear fit (zero intercept, slope = 27.9 ± 1.4 s^-1^, 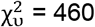) is shown in Fig. 3C (Fig. S8A shows the same data color-coded by force).

Imposing a symmetric bend angle condition where both α_1_ and α_2_ simultaneously satisfy the same constraints, α_lo_ < α_1 &_ α_2_ < α_hi_, yields a marginally higher correlation of 0.77 but a substantially worse fit 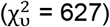. The symmetric condition is therefore not strongly supported. The data are consistent with only one segment being gently bent, while the other can adopt a range of conformations with no loss of correlation.

The correlation decreased if only one or two geometric parameters were constrained. For example, for single parameters, the correlations were: 0.46 for juxtaposition distance, 0.29 for bend angles, and 0.58 for crossing angle.

It is possible, though unlikely, that the parameters defining the preferred crossing geometry need not occur simultaneously. To test this possibility, we calculated the peak correlation between the unlinking rate and the non-simultaneous probability products (P = P (d_lo_< d < d_hi_) × P (α_lo_< α_1_ | α_2_ < α_hi_) × P (θ_lo_< θ < θ_hi_)). For this non-simultaneous probability of obtaining the preferred crossing parameters, we obtain a correlation of 0.69 and a crossing geometry with a similar bend angle (37° < α_1_ | α_2_ < 38°) as the simultaneous probability correlation approach. The higher correlation obtained with the simultaneous condition supports a model in which the preferred crossing geometry parameters likely occur simultaneously.

To confirm that the observed correlation was not an artifact of the analysis method, we performed correlation analysis with randomly permuted enzyme unlinking rates. Over 10 rounds of random permutations, the mean ± SD correlation was 0.37 ± 0.04, significantly lower than the 0.74 value obtained from unpermuted data, validating the approach.

We also performed an independent correlation analysis based only on the pairwise distances instead of calculated geometrical parameters. This alternative approach offers a qualitative perspective on the preferred crossing geometry and is consistent with the preferred crossing geometry obtained through the correlation analysis. See section S11 for details.

### The preferred crossing geometry for *M. mazei* topo VI differs markedly from that of topo IV

Applying the same joint-probability analysis to *M. mazei* topo VI experimental and simulation results from previous work (29) yielded a maximum correlation of 0.92, with preferred crossing geometry: 3.2 nm < d < 5.0 nm, 86° < α_1_ | α_2_ < 88°, 78° < < 88°. Fig. 3D shows a rendering of this preferred crossing geometry for topo VI. The corresponding scatter plot of unlinking rates as a function of the joint probability and a linear fit (zero intercept, slope = 9.4 ± 0.3 s^-1^, 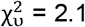) is shown in Fig. 3E (Fig. S8B shows the same scatter plot with data color-coded for positive and negative turns). The topo VI preferred geometry differs substantially from that of topo IV in both bend angle and crossing angle, despite their similar preferred crossing angles reported previously (28, 29). Random permutation of the rates gave a mean ± standard deviation correlation of 0.74 ± 0.04, again significantly lower than the unpermuted value of 0.92.

### Implications of preferred crossing geometry for chiral discrimination models

Topo IV primarily acts as a decatenase but also relaxes positive supercoils rapidly and processively (39, 42–46). *In vivo*, topo IV unlinks catenated chromosomes before cell division and relaxes positive supercoils generated during DNA replication (42–44). *In vitro*, topo IV relaxes positive supercoils at least 20-fold faster than negative supercoils(28, 39, 45, 46). This chiral discrimination has been attributed to dramatic processivity differences: topo IV is highly processive on positively supercoiled DNA and perfectly distributive on negatively supercoiled DNA (28). In contrast, although topo VI is also primarily a decatenase, it slowly and distributively relaxes both positive and negative supercoils (1, 2, 29), with a modest ~2-3 fold higher rate for positive supercoils (29).

Despite significant differences in supercoil relaxation by topo IV and top VI, the previously reported preferred crossing angles were similar: 85.5° ± 0.4° for topo IV (28) and 87.8° ± 0.4° for topo VI (29). We refer to this previous single crossing angle characterization of the crossing as the collated crossing angle, to distinguish it from the crossing angle measured in the current work. The collated crossing angle represents an average projected angle over all juxtaposition events within 10 nm at every simulation timestep (Fig. S9), convolving crossing angle with segment bending and separation (28, 29). As a result, the collated crossing angle is unable to distinguish differences in preferred segment bending or separation that contribute to differences between the two enzymes (28, 29). The collated crossing angle approach developed previously does not uniquely define the full crossing geometry of the juxtaposition and necessarily fails to capture the importance of segment bending, which differs significantly between topo IV and topo VI. The current approach of tracking the single closest pair per timestep and correlating the joint probability of multiple geometric parameters with unlinking rate isolates the catalytically relevant geometry and explicitly captures the contributions of segment bending and separation on activity while providing the complete three-dimensional geometry of the selected juxtaposition.

To determine how the differences in preferred crossing geometry between topo IV and topo VI impact supercoil relaxation, we estimated the geometric crossing parameters of a representative plectoneme. For a uniform plectoneme under 0.4 pN of force (47), we obtained the following geometric parameters: d = 11.1 nm, α = 24.4°, θ = 54.1° (Figs. 4A-B). Comparing these values with the preferred crossing geometries of topo IV and topo VI (Figs. 4C-D) reveals a clear asymmetry. The topo IV preferred crossing angle of ~55.5° is close to that of a positive plectoneme crossing and far from that of a negative plectoneme crossing; only a small fluctuation is needed for a positive supercoil crossing to satisfy the preferred geometry, whereas a much larger fluctuation is required for a negative supercoil crossing. This geometric asymmetry provides a direct mechanistic basis for the dramatic processivity difference of topo IV on positive versus negative supercoils: on positive supercoils, the preferred geometry is readily achieved, enabling processive relaxation; on negative supercoils, the required fluctuations are large and infrequent, resulting in distributive relaxation (28).

**Figure 4.**
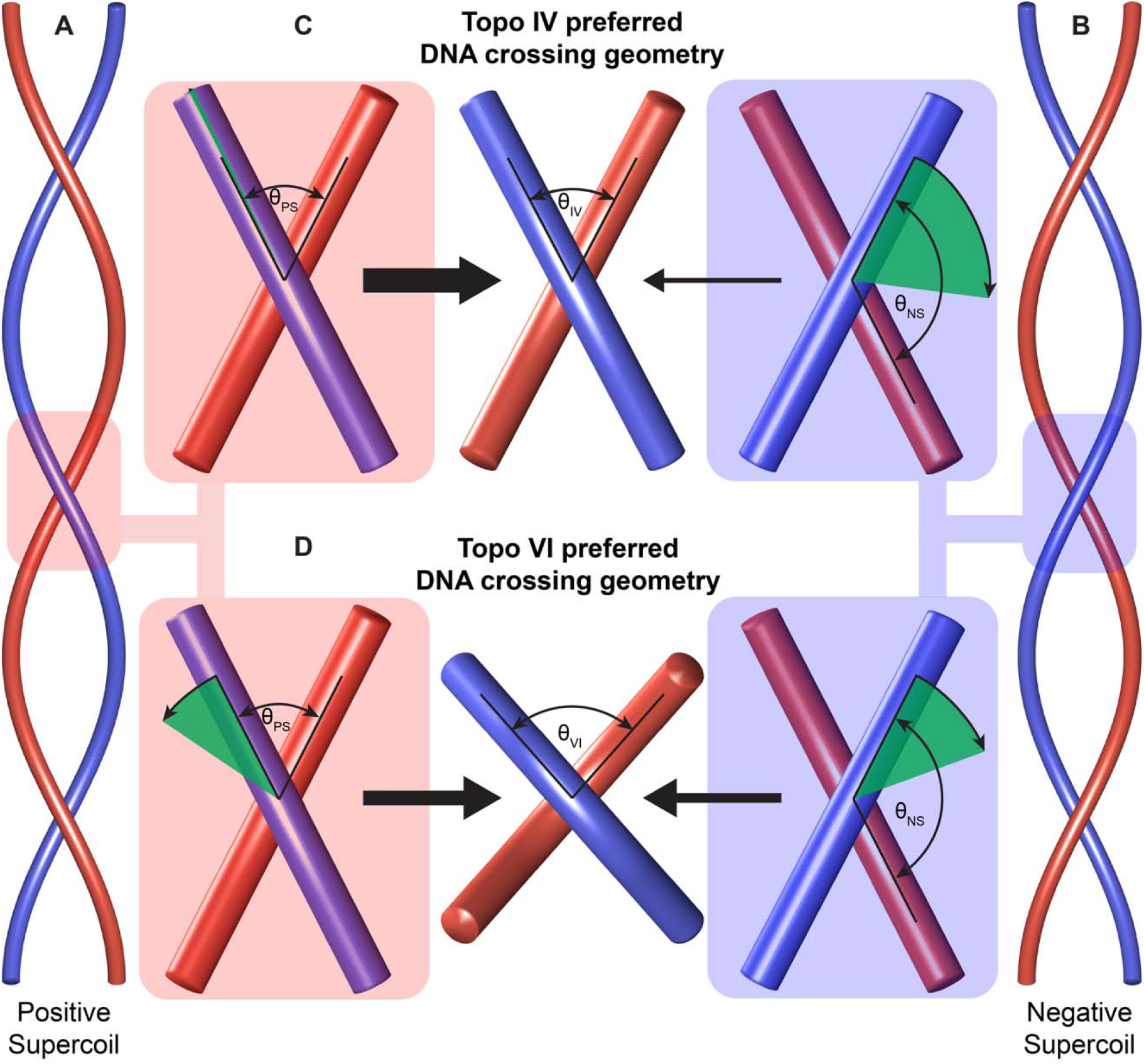
**A-B**. Positively and negatively supercoiled plectoneme under 0.4 pN of tension, respectively. **C-D. left and right figures**: a single crossing representing geometric configuration in positively and negatively supercoiled plectoneme. **C. middle figure**: Topo IV’s preferred crossing geometry. **D. middle figure**: Topo VI’s preferred crossing geometry. θ_PS_ and θ_S_ denote the crossing angle in positively and negatively supercoiled plectoneme, respectively. θ_lV_ and θ_Vl_ denote the preferred crossing angle by topo IV and topo VI, respectively. Green wedges represent the extent of relative rotation needed for a given crossing configuration to obtain the preferred crossing angle for the respective topo II.

In contrast, the topo VI preferred crossing angle range (between 78° and 88°) is poorly aligned with either positive or negative plectoneme crossings, consistent with its slow and distributive supercoil relaxation independent of supercoil chirality (29). Nonetheless, the slightly closer alignment with positive supercoils accounts for the modest ~2-fold rate preference in relaxing positive supercoiled DNA (29).

The full three-dimensional preferred geometries thus resolve the apparent paradox arising from the similar collated crossing angles of topo IV and topo VI (28, 29). Their similar θ_collated_ values obscured substantial differences in preferred bend angle and segment separation that are revealed by the joint-probability approach and that underlie the distinct behaviors of topo IV and topo VI.

### Comparison with chiral discrimination models based on crossing angle alone

The preferred crossing geometry parameters (d, α_1_, α_2_, and θ) reported here and the preferred collated crossing angle (θ_collated_) reported previously for topo IV and topo VI (28, 29) were quantified by different methods. To compare the preferred crossing geometry obtained here with the collated crossing angle reported previously, we calculate the collated crossing angle for the crossing geometry parameters obtained here for topo IV and topo VI.

For topo IV, the reported θ_collated_ = 85.5° ± 0.4° (mean ± SEM) (28). Applying the collated crossing angle calculation (Fig. S9) to the preferred crossing geometry identified here gives θ_collated_ (θ_preferred_)= 69.9° ± 16.8°, within one standard deviation of the previously reported value (28). For topo VI, the reported value of θ_collated_ = 87.8° ± 0.4° (mean ± SEM) (29), also falls within one standard deviation of the θ_collated_ (θ_preferred_) = 76.7° ± 9.6°. These results suggest that the two approaches describe the same underlying crossing geometries. However, the additional geometric parameters provided by the current approach provide a more precise and complete description of the crossing geometry, revealing important differences between topo IV and topo VI (Fig. 3B and 3D) that were not apparent from their similar collated preferred crossing angles.

### Implications of T-segment bending angle for below-equilibrium topology simplification

Our results indicate that topo IV preferentially unlinks crossings in which at least one of the segments is gently bent (Figs. 5A and 5B). We propose that the gently bent segment (red segment in Figs. 5A and 5B) is the T-segment, and the unconstrained segment (blue segment in Figs. 5A and 5B) is the G-segment. This assignment is motivated by the finding that topo IV imposes a ~126° bend upon binding the G-segment (21), significantly sharper than the ~37° bend associated with preferential unlinking. The initial bending of the G-segment does not contribute significantly to the geometric selection since the enzyme will impose the characteristic bend independent of the starting configuration. In this model, topo IV selects a T-segment that is gently bent towards the G-segment whereas the configuration of the G-segment prior to topo IV binding does not significantly impact strand passage. After binding the crossing, topo IV bends the G-segment prior to strand passage (Fig. 5D). This scenario, in which topo IIs act on a preformed juxtaposition occurring at equilibrium in which once strand is gently bent, is broadly consistent with the original HJP model (30).

**Figure 5.**
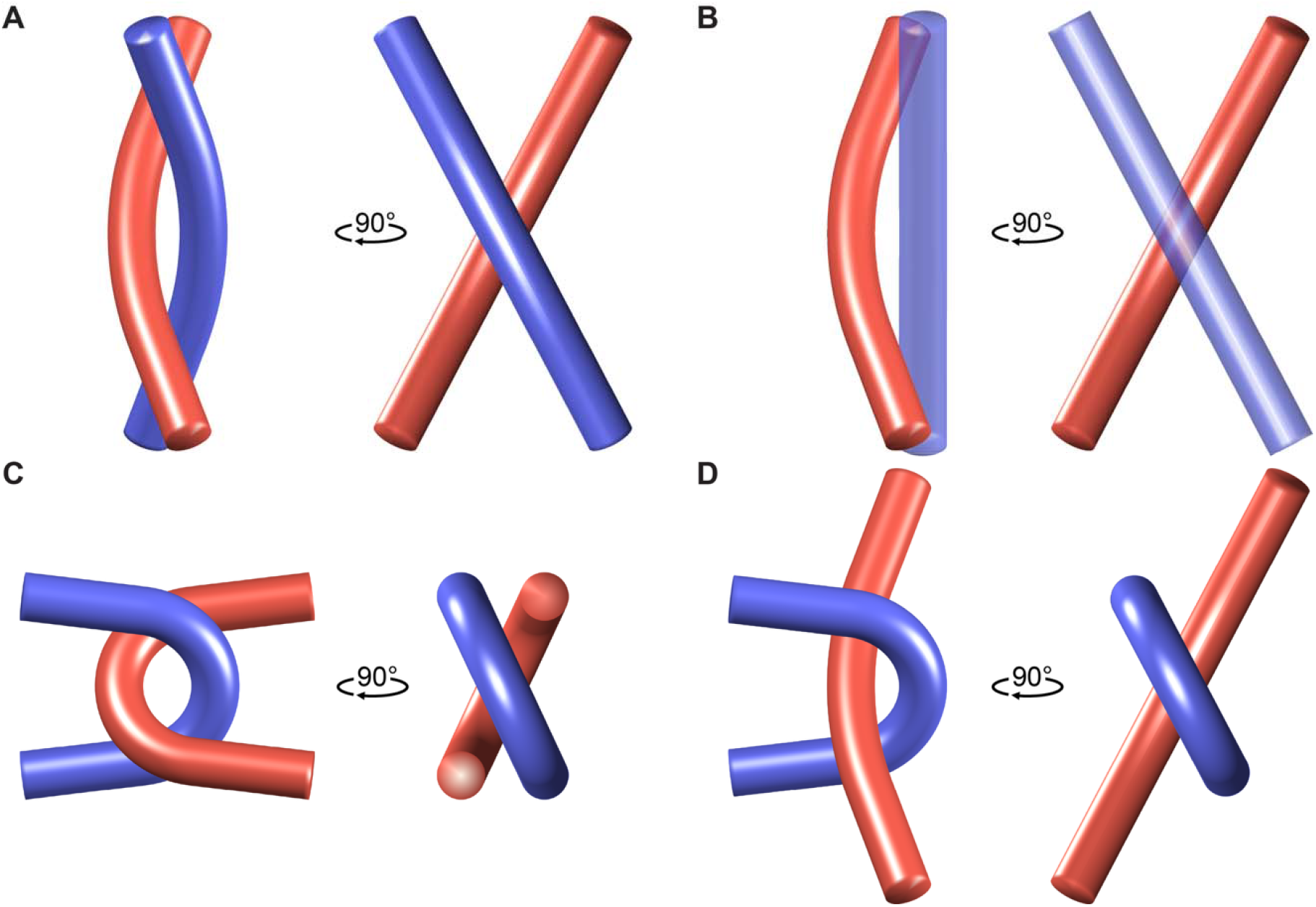
Measured and predicted preferred DNA crossing geometries for topo IV. **A.**Symmetric (fully-hooked) crossing geometry consistent with measurements: *d* = 3.75 nm, α_1_ = 37°, α_2_ = 37°, θ = 55.5°. **B**. Asymmetric (half-hooked) crossing geometry consistent with measurements: *d* = 3.75 nm, α_1_ = 37°, α_2_ = 0°, θ = 55.5°. **C**. Crossing geometry predicted by the HJP for topo IV: *d* == 5 nm, α_1_ = 165°, α_2_ = 165°, θ = 50°. **D**. Crossing geometry illustrating the sharp bend imposed by topo IV on the G-segment (blue) and gently bent T-segment (red): *d* = 3.75 nm, α_1_ = 37°, α_2_ 165°, θ = 55.5°. For each figure, the illustration on the left shows the front view and, on the right, shows the side view of the crossing.

Subsequent extensions of the HJP model demonstrated that preferential unlinking of sharply hooked juxtapositions can produce below-equilibrium topology simplification (31–36). The preferred crossing geometry for topo IV offers a direct test of the main prediction of these extended HJP models. Below-equilibrium relaxation of circular DNA is quantified by R_Lk_, the ratio of the variances of equilibrium to below-equilibrium steady-state topoisomer distributions quantified as linking number (Lk) (R_Lk_=<Lk^2^ _eq_>/<Lk^2^ _non-eq_ >) (15). Using the quantitative modeling relating the geometric parameters (α_1_, α_2_, θ, d) to the degree of below-equilibrium topology simplification, we calculated the geometry predicted to obtain the reduction in supercoiled DNA variance R_Lk_ = 1.77 for *E. coli* topo IV (15), and compared it with the preferred crossing geometry (Fig. 5C). Whereas the predicted and measured *d* and θvalues are comparable, the HJP model predicts a sharply and symmetrically bent crossing (α_1_ =α_2_ ≈ 165°) that is inconsistent with the measured geometry (α_1_| α_2_ ≈ 37°). Bending angles of ~165° were not observed in any of the simulated trajectories (Figs. 2E and 3A).

Since the HJP model for below-equilibrium topology simplification is based on the assumption that topo IIs recognize a pre-existing DNA juxtaposition geometry (36), we compare the bending energies of the predicted and calculated crossing geometries. The bending energy of the sharp (165°) bending of both segments is ~33 k_B_T, making such a configuration extremely improbable. Conversely, the bending energy for a bend of 37° is ~1.7 k_B_T (See section S12 for details), and the relative probability of the highly bent crossing occurring compared to the gently bent crossing is extremely low (~e^−31^ ≈ 3×10^-14^). Sharply hooked juxtapositions are therefore not viable substrates for geometric selection as postulated by the extended HJP model for non-equilibrium topology simplification.

We consider a slightly relaxed version of the HJP model in which topo II is stably bound at the G-segment and selects a T-segment with the appropriate geometry for strand passage. This scenario is consistent with the data indicating that strand passage is dictated by the bending of only one of the DNA segments at the crossing, the T-segment. Although the modeling did not explicitly consider the possibility of one segment being sharply bent to the extent expected for G-segment bending, we note that the bending of the two segments is uncorrelated (Fig. S3B). Thus, to a first approximation, a sharp bend imposed on the G-segment would not impact the bending of the T-segment. In this scenario, our analysis reports on the bending of the T-segment required for strand passage, which we find is gently bent. Summarizing the interpretation of the results in the context of the stringent or relaxed version of the HJP model, we find that topo IV selects a crossing in which the T-segment is gently bent, independent of the bending of the G-segment.

The gentle T-segment bending identified here, which is a key qualitative insight of the original HJP model, contributes to topology simplification within the extended HJP model, predicting R_Lk_ ~ 1.33. The full measured value of R_Lk_ = 1.77 likely requires additional mechanisms, which may vary among topo II enzymes in a manner consistent with the range of R_Lk_ values reported by Rybenkov et al (15).

Recent cryo-EM structures of *E. coli* DNA gyrase bound to supercoiled DNA provide complementary support for our findings (24, 25). One structure shows a crossing with a gently (~20°) bent T-segment (24), and G- and T-segments juxtaposed at ~60°, similar to our findings for *E. coli* topo IV. A second structure supports gentle T-segment bending in the context of chirally-wrapped DNA (25). Direct comparison with our results is limited by the functional differences between topo IV (primarily decatenation) and gyrase (primarily relaxing positive supercoils and introducing negative supercoils), and by the possibility that the cryo-EM structures capture a subset of the dynamic conformations sampled in solution. Nevertheless, the agreement between the static gyrase structures and the kinetically selected geometry reported here is notable. Obtaining the kinetically selected crossing geometries for other type IIA and IIB topos will establish the degree of conservation and functional variation among topo IIs.

## Discussion

We developed a kinetics-based approach to determine the complete three dimensional geometry of the G- and T-segments selected by topo II for strand passage. The approach combines single-molecule measurements of topo II unlinking rates over a range of experimentally imposed DNA crossings with Brownian dynamics simulations to obtain the geometric parameter distributions of the crossings. Maximizing the correlation between measured unlinking rates and the joint probability of obtaining specific geometric parameter combinations identifies the full three-dimensional G- and T-segment geometry that is kinetically selected for strand passage.

This kinetic approach reports the juxtaposition geometry that is selected for activity, which may differ from the geometry captured in static structures of topo II with both segments bound. Static structures provide snapshots that may or may not represent the preferred pre-binding geometry, whereas the kinetic approach directly reports the geometry selected for strand passage.

For *E. coli* topo IV, the preferred crossing geometry is defined by: d = 3.75 ± 1.5 nm, α = 37°± 2°, θ = 55.5° ± 0.5°, where d is the distance between the juxtaposed segments, α is the bending of the juxtaposed T-segment, and θ is the angle between the juxtaposed G- and T-segments. The two DNA segments are in close proximity, acutely crossed, and one segment, assigned as the T-segment, is gently bent, while the second segment is largely unconstrained. For *M. mazei* topo VI the preferred crossing geometry is defined by: d = 4.1 ± 0.9 nm, = 87°± 1°, = 83° ± 5°. The two DNA segments are in close proximity, are acutely (and near-orthogonally) crossed, and the assigned T-segment is near-orthogonally bent, while there is little constraint on the G-segment. In comparison with topo IV, the topo VI preferred geometry has a similar juxtaposition distance, but substantially larger bending and crossing angles.

These preferred crossing geometries extend and refine the chiral discrimination model in two important ways. First, the full geometric description reveals significant differences between topo IV and topo VI that were obscured when only the single collated crossing angle was considered (28, 29). Second, the geometry of the preferred crossing provides a mechanistic basis for the contrast between processive relaxation of positive supercoils by topo IV and distributive relaxation by topo VI. The topo IV preferred crossing angle of ~55.5° closely matches that of a positive plectoneme crossing, requiring only small fluctuations for the preferred geometry to be achieved and enabling processive relaxation. The much larger fluctuations required for negative supercoils result in distributive relaxation. The topo VI preferred crossing angle of 78°–88° is poorly matched by crossings in either positive or negative plectonemes, explaining its slow and distributive supercoil relaxation regardless of chirality, with a modest preference for positive supercoils.

The preferred crossing geometry for topo IV is inconsistent with the sharply and symmetrically hooked juxtaposition predicted by extended HJP models as the basis for below-equilibrium topology simplification by topo IV. The predicted bend angles of ~165° in both segments are not observed in any simulated trajectory and carry an energetic cost (~33 k_B_T) that makes them overwhelmingly improbable. The gentle T-segment bending that is observed — a qualitative feature of the original HJP model — does contribute to topology simplification, yielding a predicted simplification factor R_Lk_ ~ 1.33, but falls short of the measured value of 1.77 (15). The remaining discrepancy implies that additional, as yet unidentified mechanisms contribute to below-equilibrium simplification, potentially varying among topo II enzymes in ways that account for the range of R_Lk_ values reported across the type IIA family (15).

The approach developed here is general. Correlating strand-passage kinetics with the joint probability of crossing geometry parameters can be applied to any enzyme that resolves a DNA synapse, providing a measure of the kinetically selected synapse geometry that complements and extends structural approaches. Applied to topo IV and topo VI, it has revealed distinct preferred crossing geometries that resolve a paradox, provide a geometric basis for chiral discrimination and processivity differences, and set quantitative constraints on models of below-equilibrium topology simplification.

## Materials and Methods

### Topoisomerase IV unlinking substrate

Linear 3.6 kb dsDNA labeled with biotin or digoxigenin at each 5□ end was prepared by PCR using pET28b (EMD Science) as a template and biotin (bio)-labeled primer (5□ Bio-Bio-ctgttcatccgcGTCCAGCTCGTTG) and digoxigenin (dig)-labeled primer (5□-Dig-Dig-Dig-GGACCTGCTTTCGTTGGCGTAATGGCTGGCCTGTTG) (Eurofins MWG Operon). The PCR product was purified with a QIAquick PCR purification kit (Qiagen).

### *E. coli* Topo IV

The ParC and ParE subunits of *E. coli* topo IV were purified as previously described (21, 48). Equimolar amounts of ParC and ParE were mixed to make a stock of 14 μM topo IV heterotetramer.

### *M. mazei* Topo VI

Topo VI purification and activity measurements are described in our previous work (29).

### Single-molecule DNA crossing unlinking assay

To prepare double DNA tethers, 1 µl of 0.15 nM 3.6 kb dsDNA was incubated with 1 µl of magnetic beads (1% w/v, MyOne, Invitrogen) in 40 µl of wash buffer (WB: 1×Phosphate-buffered saline (PBS), 0.3% w/v bovine serum albumin (BSA), and 0.04% Tween-20) overnight at 4°C at a ~3:1 DNA to bead ratio. The sample cell, composed of two coverslips attached together with ~50 µm double-sided adhesive (8132LE, 3M), was incubated with 40 µl of 5 µg/ml anti-digoxigenin in 1x PBS overnight at 4°C, followed by a wash with 600 µl of WB. The DNA-bead mixture was incubated in the anti-digoxigenin-coated sample cell for 30 min before washing with 2 ml of WB to remove unbound DNA and beads. The magnetic tweezers instrumentation has been previously described (49–51).

Double-DNA tethered beads were identified by rotating the magnet assembly. In contrast to a bead tethered by a single DNA, the DNA extension of a double-DNA tethered bead decreases symmetrically upon both right-handed and left-handed bead rotation (52). Topo IV activity buffer (25 mM Tris HCl, pH 7.5, 100 mM potassium glutamate, 10 mM magnesium chloride, 1 mM dithiothreitol, 0.3% w/v BSA, and 0.04% v/v Tween-20) containing 2 nM *E. coli* topo IV (above the dissociation constant, K_d_ of ~0.2 nM measured under similar conditions) (41) and 1 mM ATP was introduced once double-tethered beads were found and their crossing geometries (spacing between DNA molecules and verification of parallel tether geometry described below) were characterized.

The kinetic measurements report the rate-determining step or steps of the unlinking reaction. Working at protein concentrations well above K_d_ and with saturating ATP ensures that neither protein binding nor ATP binding is rate-determining. Nonetheless, a priori, we do not know if the rate-determining step in the strand-passage cycle depends on the crossing geometry.

Experimental verification that the unlinking rate depends upon the imposed DNA geometry confirms it is rate-determining (Fig. 1). Furthermore, the single exponential distribution of unlinking times indicates only one irreversible rate-determining step in the unlinking reaction.

Unlinking measurements were performed by applying crossings of 0.7 to 1 turn after each unlinking event, which was identified by the bead height increase. This process was repeated for 30 or more unlinking events per crossing. Bead height traces were analyzed using a custom step-finding algorithm to identify the waiting time, T_wait_ (the interval between rotating the bead to generate the DNA crossing and it being unlinked by the topoisomerase) (38). T_wait_ values were binned and fitted with a single exponential to obtain the unlinking rate for each imposed rotation.

The current study focuses on the effect of crossing geometry using shorter DNA molecules (3.6 kb) than what was previously used (5 kb DNA) (38), which permits accessing a broader range of crossing geometries and, in particular, a broader range of bend angle (α) (Fig. S10).

### DNA crossing geometry determination

The geometry of the two DNA segments tethering the bead to the surface was determined by measuring the extension, Z(R), as a function of the number of turns R. The region from −0.5 to 0.5 turns was fitted with Eq. 2 to obtain the distance between the two DNA molecules (S) and the DNA extension at R = 0 (Z_0_) (Figs. 1B and 1C):

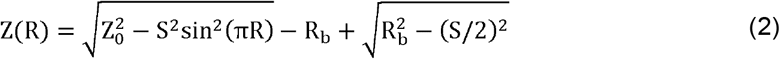

where Z(R) is the distance between the glass surface and the bottom of the bead, and R_b_ is the bead radius (0.5 μm) (28, 38). In Eq (1), the first term on the right describes the extension of a parallel tether as a function of rotation, and the other two terms are constant corrections that account for the change in the measured extension due to the curvature of the bead (28, 38). Tethered beads were selected for unlinking measurements if the DNA extension at zero rotation (Z_0_) was within 10% of the predicted extension at the applied force, eliminating non-parallel and multi-tether geometries. Beads tethered by more than two DNA molecules were additionally excluded by rejecting tethers with asymmetric extension versus rotation curves.

### Coarse-grained modeling of DNA crossings

We performed coarse-grained Brownian dynamics simulations of each experimental DNA crossing geometry with the experimental values of force, DNA separation, and length, and rotation using a standard coarse-grained model (53–57) to obtain the local crossing geometry. The simulation model is summarized below; additional details are provided in our previous work (29).

In the simulations, two linear polymers consisting of charged monomers are arranged in a parallel configuration at a fixed separation, mimicking the double tether in the experiment. One end of the double-molecule is rotated about its long axis while the other end is kept fixed. This rotation mimics the rotation of the DNAs attached to the magnetic bead in the experiments. Each polymer segment comprises 486 monomer beads of 2.5 nm diameter, σ, corresponding to a 3600 bp long dsDNA segment, and hence each monomer bead represents 7.4 bp of dsDNA with a −2.96 e charge (53). Simulation parameters are reported in the reduced Lennard–Jones (LJ) units. The equivalent SI units are provided in Table S1.

The total interaction potential between monomers is expressed as sum of four potentials:

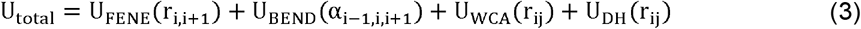

The four terms describe, respectively: a finitely extensible non-linear elastic (FENE) bond potential between neighboring monomers; a bending rigidity term; excluded volume interactions; and electrostatic interactions.

The U_FENE_ potential is :

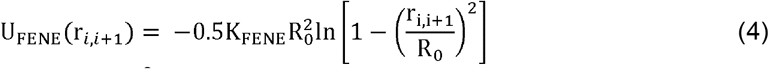

where the constants *K*_FENE_ = 30k_B_T/σ^2^ and R_0_ = 1.6σ define the bond energy and maximum bond length, respectively. Here, σ represents length in reduced units (~2.5 nm in real units) k_B_T is thermal energy.

Bending rigidity is modeled with the Kratky–Porod potential:

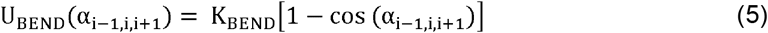

Here, K_BEND_ 20k_B_T is the bending rigidity and α_i−,i,i+1_ is the deflection angle made by three adjacent monomers (with respect to a straight line) on the polymer chain. The bending rigidity sets the persistence length, l_p_ = 20σ = 50 nm.

Excluded volume interactions are described by the Weeks–Chandler–Andersen (WCA) potential:

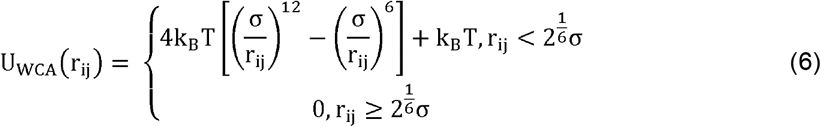

Electrostatic interactions between monomers are described by the Debye-Hückel potential:

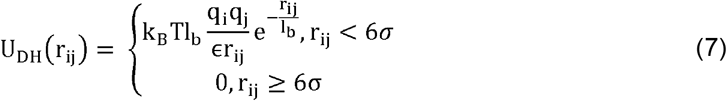

where l_b_ = (3.66σ) ^-1^ is the Bjerrum length, q_i_ 644 is the charge on particle i, and, ϵ = 1.6 is the dielectric constant, chosen to match the experimental salt concentration.

All simulations were performed using the Large-scale Atomic/Molecular Massively Parallel Simulator (LAMMPS) (58) in an NVT (a constant number of particles, volume, and temperature) ensemble with a Langevin thermostat, timestep Δt = 0.01τ (where τ is time in the reduced LJ units), and total duration T_total_ = 2×10^7^τ. The data were sampled every 250τ. The sampled crossings represent the equilibrium fluctuations prior to enzyme binding; modeling the DNA structure after topo II binding is beyond the scope of the current work.

### Identifying a crossing and defining participating segments

At each sampled timestep, the crossing is defined as the point of closest approach between the two DNA molecules. Two beads (one bead from each segment) with the minimum Euclidean distance between the two juxtaposed DNA molecules define the central beads of their respective segments. The juxtaposed segments comprise the two neighboring beads on either side of the central beads, giving a 5-bead arc with a contour length of 12.5 nm. All subsequent geometric calculations are performed on these segment pairs.

### Calculating the geometric parameters of a crossing

Four parameters characterize the crossing geometry at each timestep (Fig. S2 C-D). The juxtaposition distance, d, (Fig. S2C) is defined as:

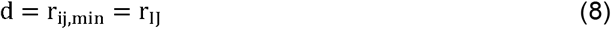

where the I and J beads are the central beads of the two juxtaposed segments. The crossing angle, θ (Fig. S2D), is defined as:

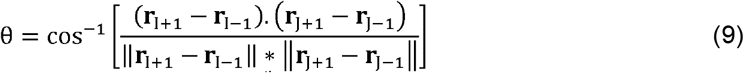

where **r**_i_ is the position of the center of mass of the i^th^ bead. The bend angles, α_1_ and α_2_ (Fig. S2C), are defined as:

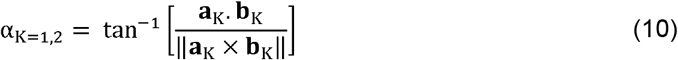

where

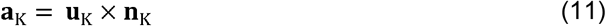

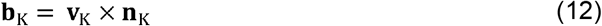

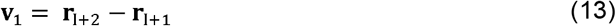

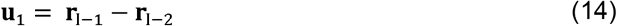

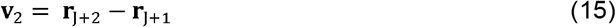

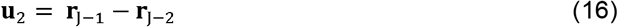

and **n**_K_ is the normal vector to the best-fit plane of each segment obtained by singular value decomposition.

In brief, α is the angle between the projections of a segment’s first and last bond vectors onto this plane. Note that the value of α follows the intuitive sense of bending i.e., for a straight segment, α=0, and the value of α increases as the segment bending increases (Fig. S2C).

The crossing angle (θ) defined here differs from the collated crossing angle, θ_collated_, reported in prior work (28, 29). Here, a unique crossing is defined per timestep as the single closest-approach pair, and a single value of θis calculated for that pair alone. In prior work, all crossings within a juxtaposition distance of 10 nm were considered at each simulation timestep and θ_collated_ was their mean (28, 29). The equivalence of the two approaches is demonstrated in Results and Discussion and Fig. S9 provides a schematic comparison.

### Visualization

We used OVITO (59) to visualize the simulated trajectories of the coarse-grained DNA beads. We used MATLAB to perform post-processing calculations to obtain the crossing parameters, and to generate renderings of crossing and plectoneme geometries.

## Acknowledgments

This work utilized the computational resources of the NIH HPC Biowulf cluster (http://hpc.nih.gov). This work was supported by the Intramural Research Program, National Heart, Lung, and Blood Institute, National Institutes of Health. We thank Shannon McKie for providing *M. mazei* Topoisomerase VI data.

## Supporting Information

### S1. Topo IV unlinking rate as a function of bead rotation

**Figure S1:**
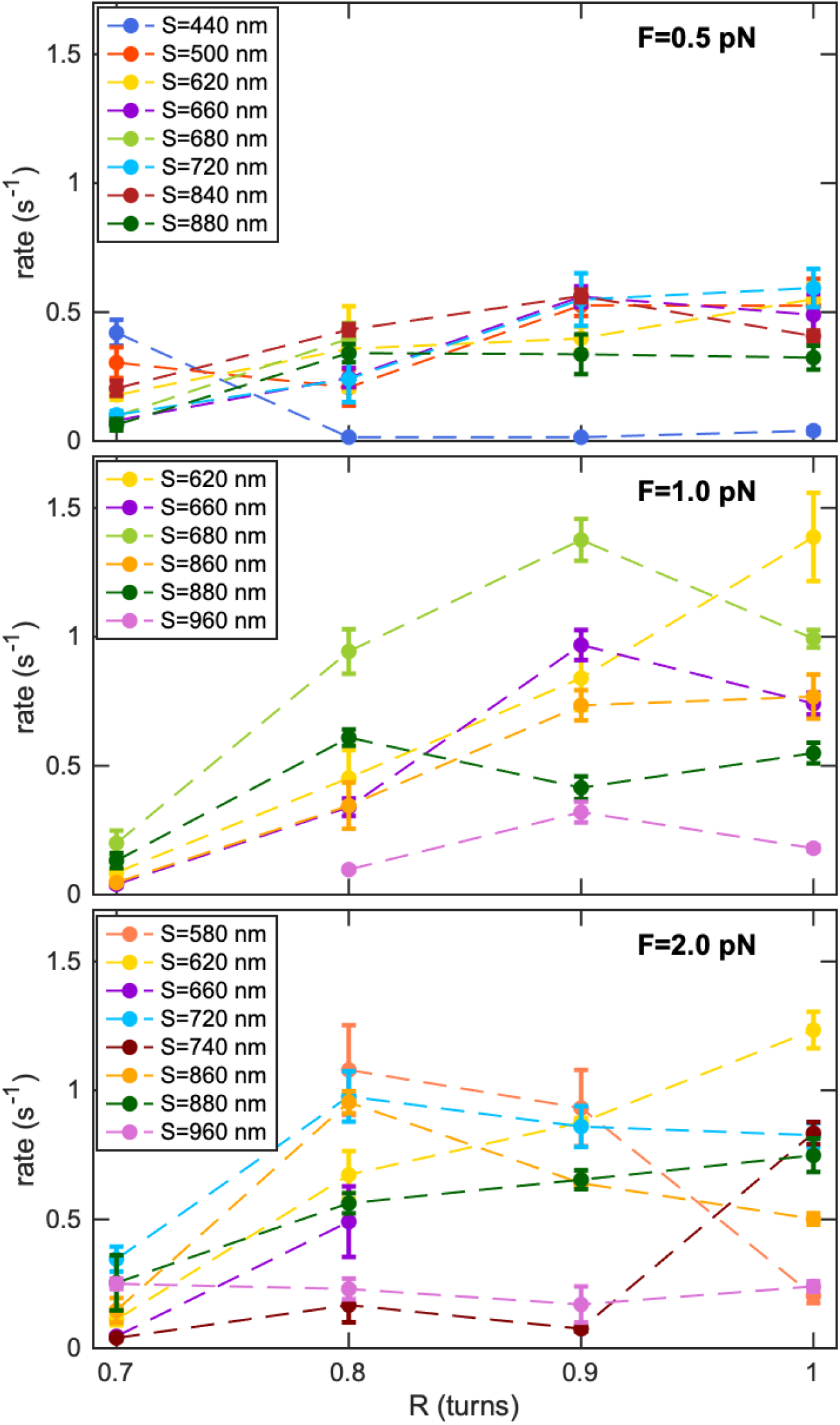
Topo IV unlinking rate as a function of bead rotation, R, for 12 different segment spacings (S) and three different forces (F). The unlinking rates measured are color-coded based on the segment spacing (S = 440 nm: Royal blue; 500 nm: Orange-red; 580 nm: Coral; 620nm: Gold; 660 nm: Dark violet; 680 nm: Yellow-green; 720 nm: Deep sky blue; 740 nm: Maroon; 840 nm: Firebrick; 860 nm: Orange; 880 nm: Dark green; 960 nm: Orchid) for three different forces (F=0.5: Top panel; 1 pN: Middle panel; 2 pN: Bottom panel). Dashed lines between points are included to guide the eye. Error bars correspond to the standard error of the mean.

### S2. Schematic of DNA crossing and parameter definitions

**Figure S2:**
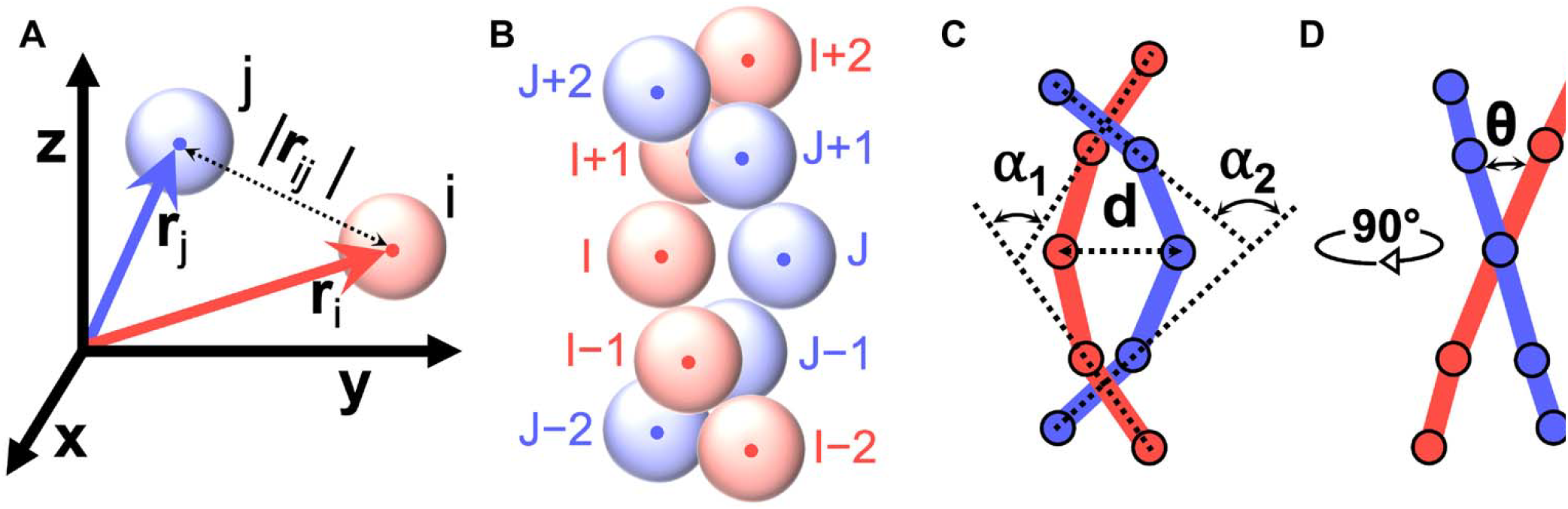
Schematic of DNA crossing and parameter definitions. **A**. Representative position vectors of monomer beads. **B**. Schematic of a crossing consisting of 5-monomer-bead segments from each dsDNA. **C**. Simplified schematic of a crossing with the ball-and-stick model (representing particle-and-bond) for easier visualization. The juxtaposition distance (*d*) is defined as the distance at the closest approach, i.e., central beads of the pair. The bend angles (*α*_1_, *α*_2_) are defined as the angle between the first and last bonds of the respective segments. **D**. Side view of the previous schematic. The crossing angle (*θ*) is defined as the angle between the planes defined by the two segments.

### S3. Similarity of the probability distributions of α_1_ and α_2_

**Figure S3:**
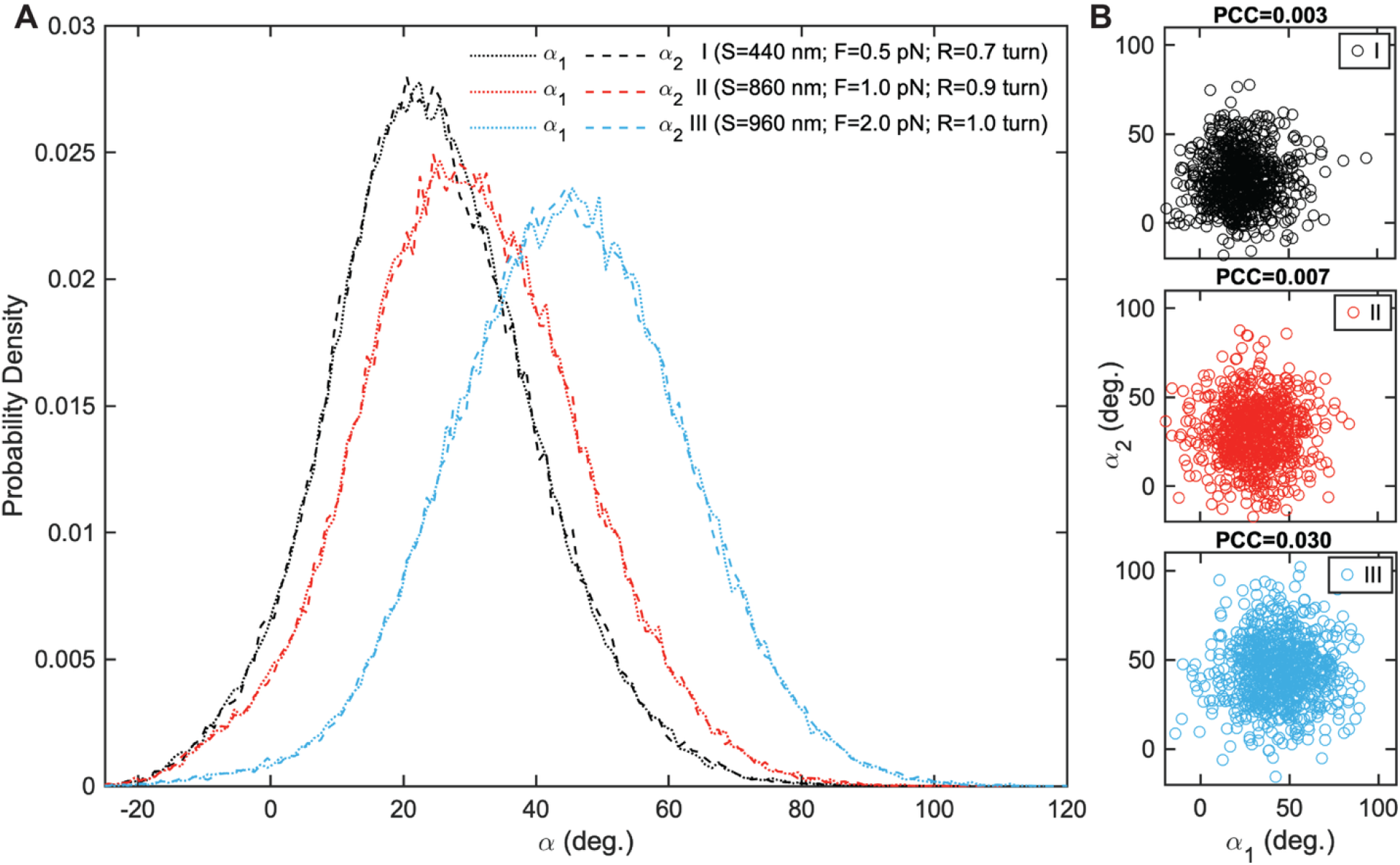
Similarity of representative probability distributions of α_1_ and α_2_ **A**. Probability distributions of α_1_ (short dashed lines) and α_2_ (long dashed lines) at three different levels of Hookedness. **B**. Scatter plots of α_2_ as a function of α_1_ at three different levels of Hookedness. density Each dot represents α for every 100^th^ timestep from a simulation for a given configuration. The Pearson correlation coefficient (PCC) is included at the top of each scatter plot. The histograms and scatter plots are color-coded for each tuple ({S, F, R} = {440 nm, 0.5 pN, 0.7 turn}: Black; {860 nm, 1 pN, 0.9 turn}: Red; {960 nm, 2 pN, 1 turn}: Blue).

### S4. The mean values of crossing parameters at F = 0.5 pN

**Figure S4:**
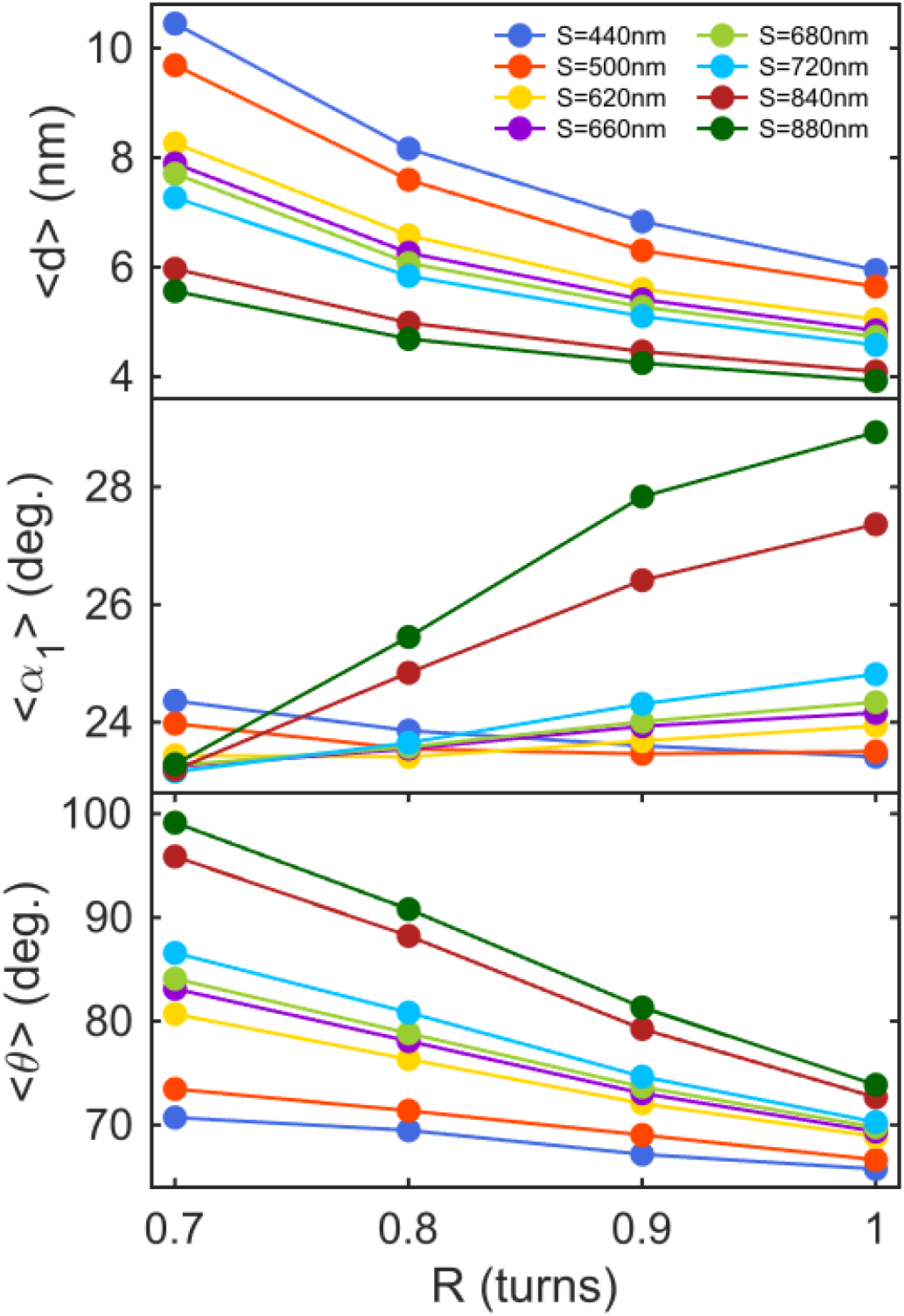
The mean values of crossing parameters, <*d*>, <_1_>, < >, as a function of bead rotation (R) from simulations conducted at 0.5 pN force for different segment separation (S) values. The mean values from simulations performed at different segment separations are color-coded (S = 440 nm: Royal blue; S = 500nm: Orange red; 620 nm: Gold; 660 nm: Dark violet; 680 nm: Yellow green; 720 nm: Deep sky blue; 840 nm: Firebrick; 880 nm: Dark green).

### S5. The mean values of crossing parameters at F = 1 pN

**Figure S5:**
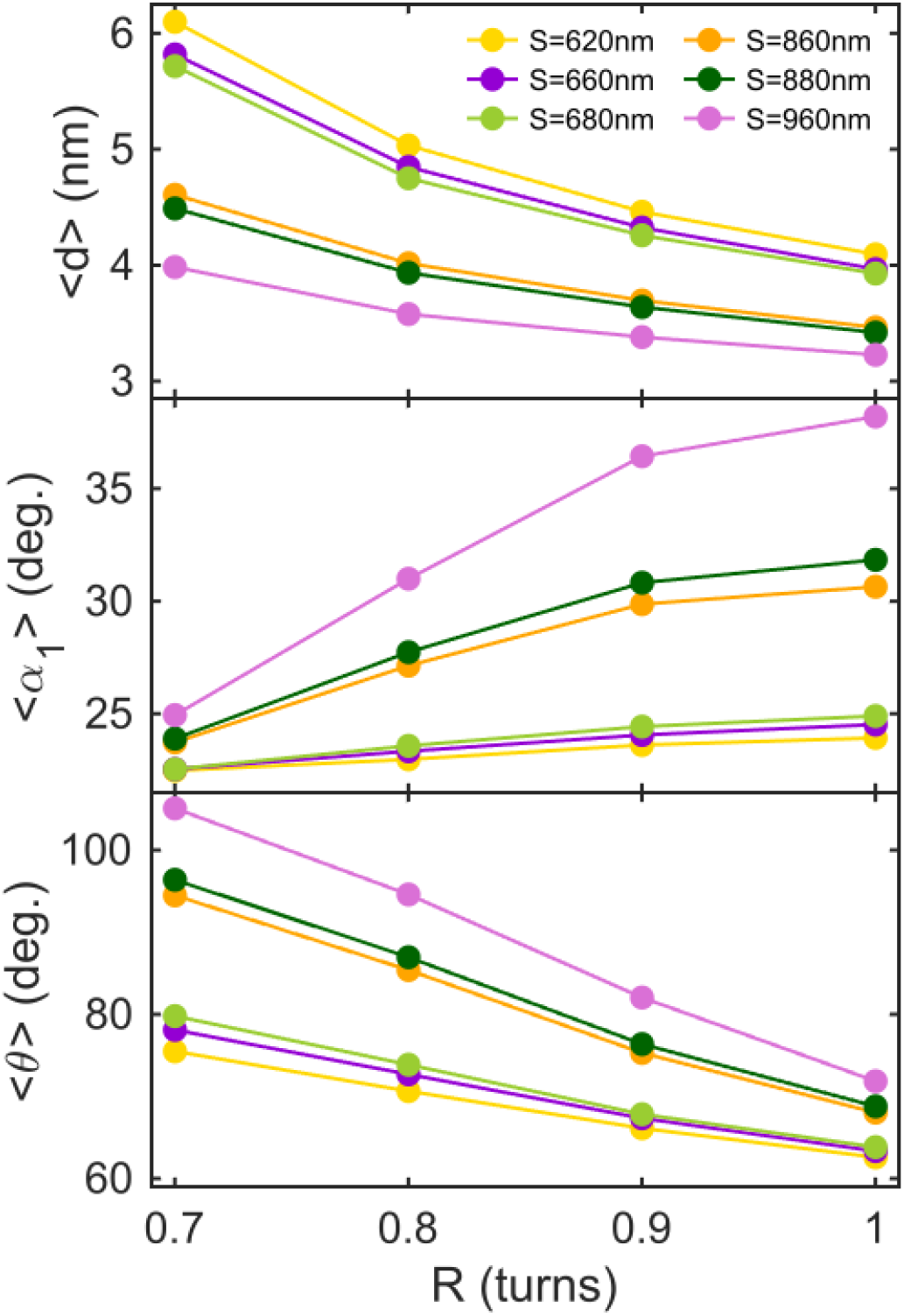
Mean values of geometric crossing parameters, <*d*>, < _1_>, < >, as a function of bead rotation (R) from simulations conducted at F = 1 pN (S = 620nm: Gold; 660 nm: Dark violet; 680 nm: Yellow green; 860 nm: Orange; 880 nm: Dark green; 960 nm: Orchid).

### S6. The mean values of crossing parameters at F = 2 pN

**Figure S6:**
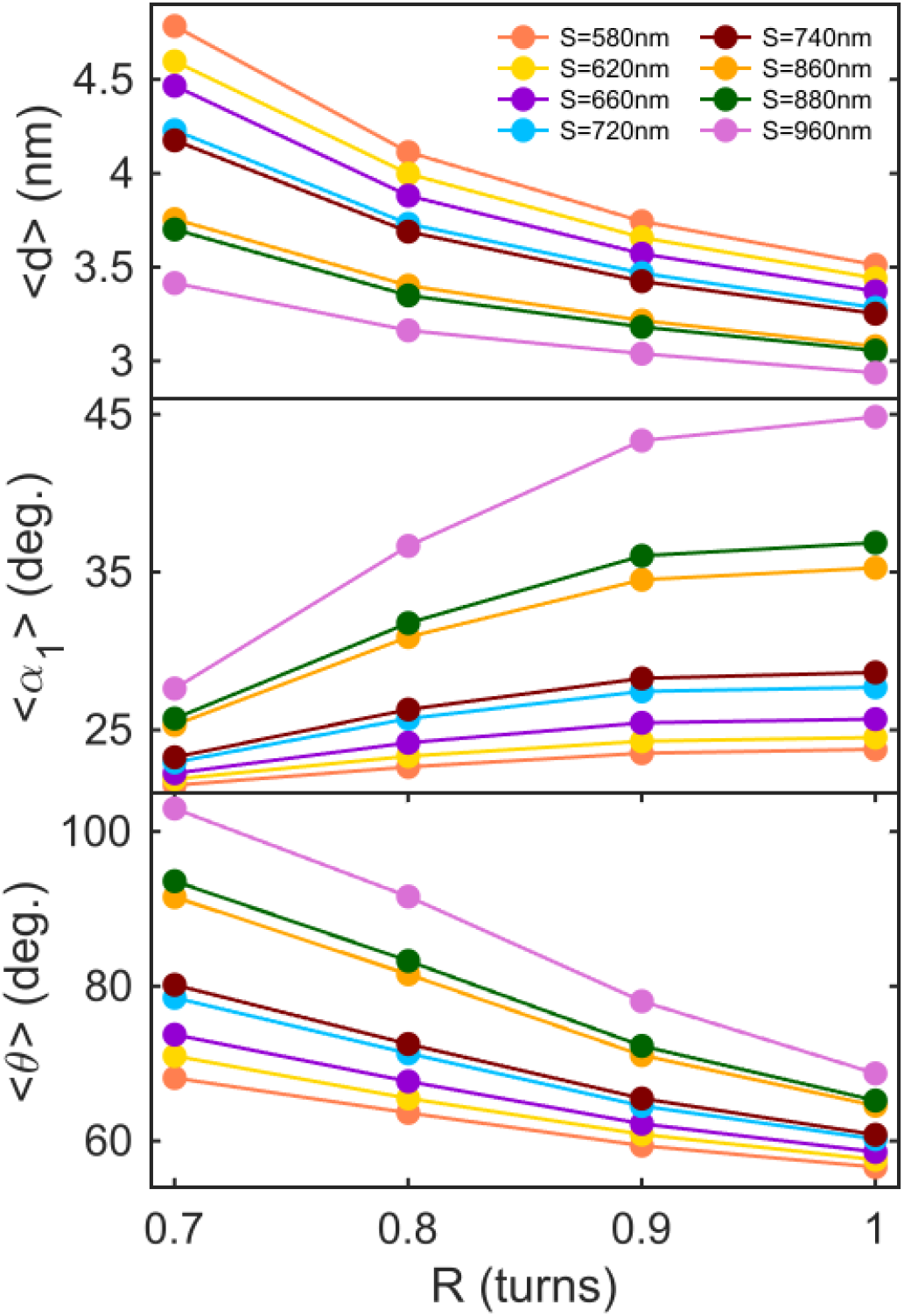
The mean values of crossing parameters, <*d*>, < _1_>, < >, as a function of bead rotation (R) from simulations conducted at 2 pN force for different segment separation (S) values. The mean values from simulations performed at different segment separations are color-coded (S = 580nm: Coral; 620 nm: Gold; 660 nm: Dark violet; 720 nm: Deep sky blue; 740 nm: Maroon; 860 nm: Orange; 880 nm: Dark green; 960 nm: Orchid).

### S7. Enzyme unlinking rate as a function of mean values of crossing parameters

**Figure S7:**
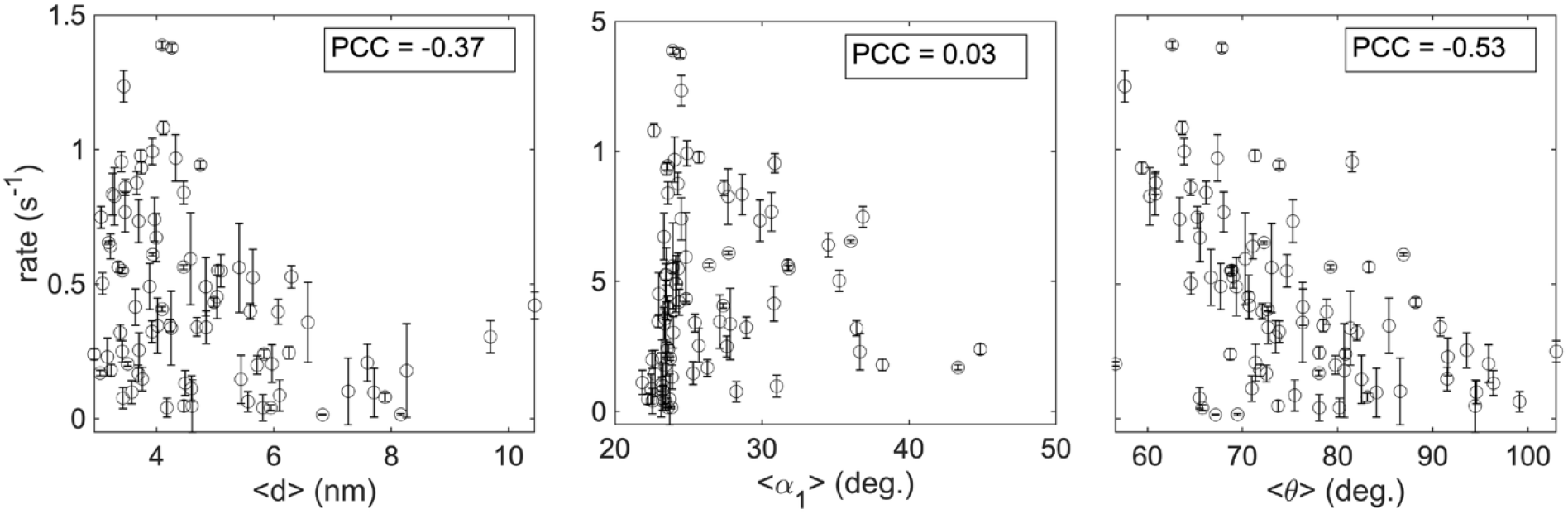
Scatter plot of enzyme unlinking rate as a function of computed mean values of crossing parameters, <*d*>, < α_1_>, and < θ > along with the computed Pearson correlation coefficients (PCC).

### S8. Enzyme unlinking rate as a function of computed joint probability for highest correlated geometric parameters

**Figure S8:**
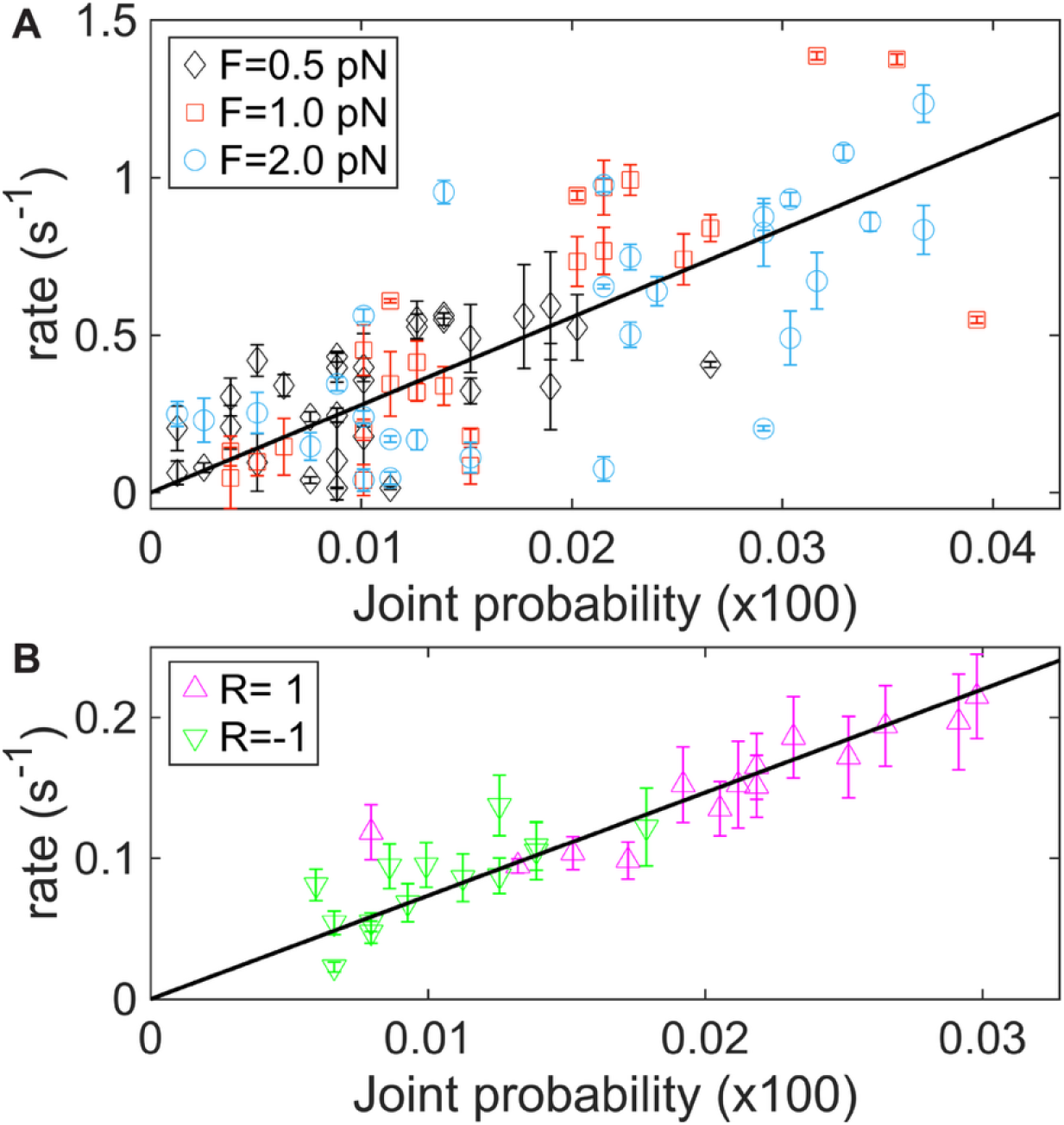
Scatter plots of enzyme unlinking rate as a function of computed joint probability with geometric parameters maximizing the correlation. **A**. Topo IV unlinking rate plotted as a function of the joint probability. The line shows a linear fit (slope = 27.9 ± 1.4 S^-1^, with 0 intercept, and educed chi-square, reduced chi-square, 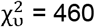). To identify possible effects of force on the measured unlinking rate independent of the geometry, we computed the reduced chi-squared for the data at each force independently: 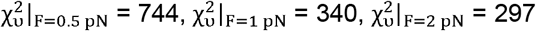. The decrease in the reduced chi-squared for with increasing force, and the lack of a systematic shift in the unlinking rates measured at different forces suggest that the rates do not decrease dramatically over the force range (0.5-2 pN). The unlinking rates for different forces are color- and symbol-coded (F=0.5 pN: Black diamond; 1 pN: Red rectangle; 2 pN: Blue circles). **B**. Topo VI unlinking rate (from previous work (1)) plotted as a function of the joint probability. The line shows a linear fit (with 0 intercept, slope = 9.4 ± 0.3 S^-1^, and reduced chi-square, 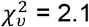). The unlinking rates for different crossing signs (1 full positive or negative turn) are color-coded (R=1: Magenta upward triangle; R=-1: Green downward triangle). Error bars correspond to the standard error of the mean.

### S9. Calculation of the collated crossing angle for all the pairs with a juxtaposition distance below a threshold value

**Figure S9:**
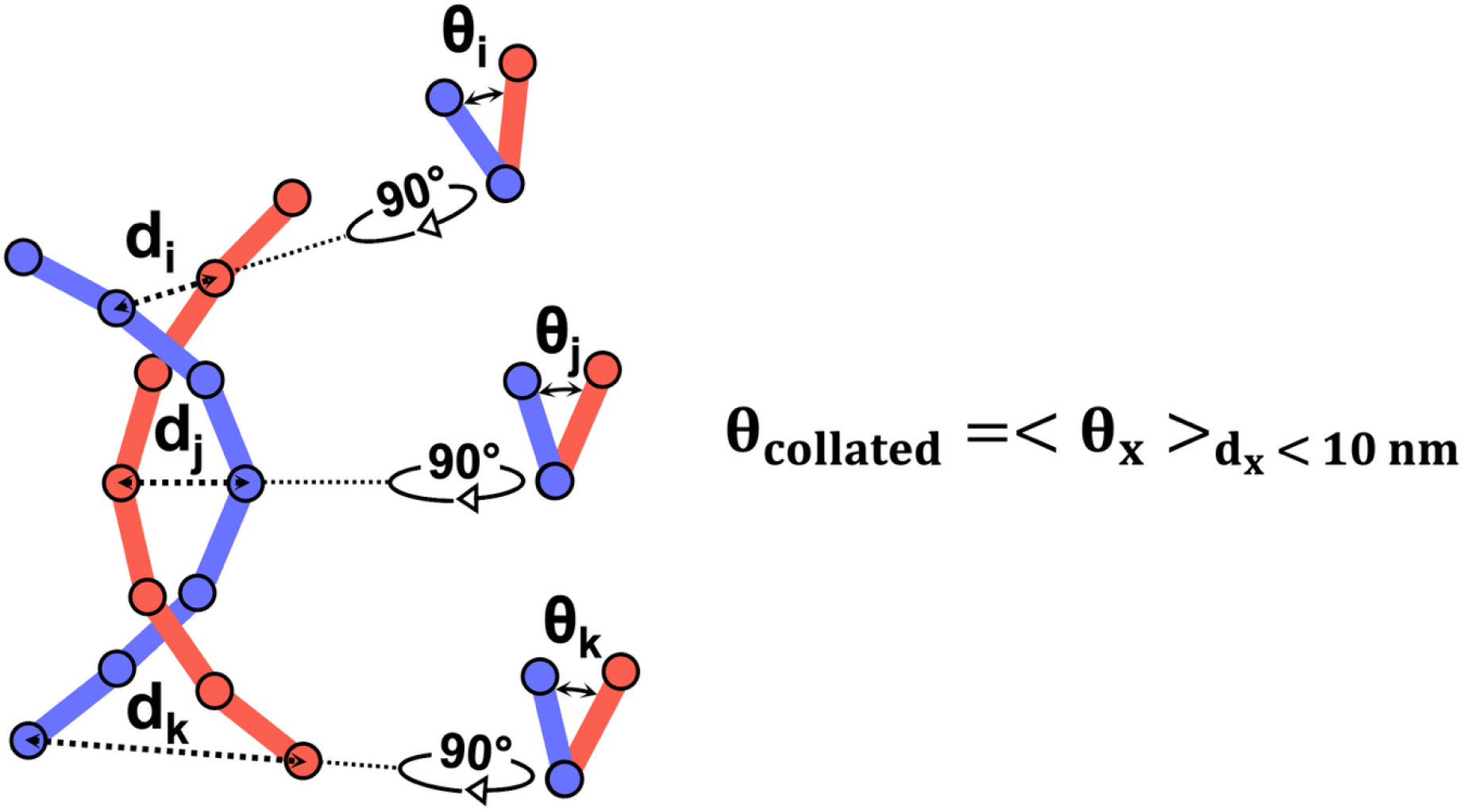
Schematic describing the calculation of collated DNA crossing angle in previous work (1, 2). To avoid confusion, we denote the crossing angle defined in prior work as. At each simulation timestep, all possible crossings with a juxtaposition distance below a threshold value, 10 nm, were considered and the crossing angle was calculated for all such crossings (1, 2). The reported crossing angle () in the previous work is the mean of individual crossing angle values measured at a given timestep (1, 2).

### S10. Comparison of crossing geometries achieved in single DNA-crossings for different sizes of DNA

**Figure S10:**
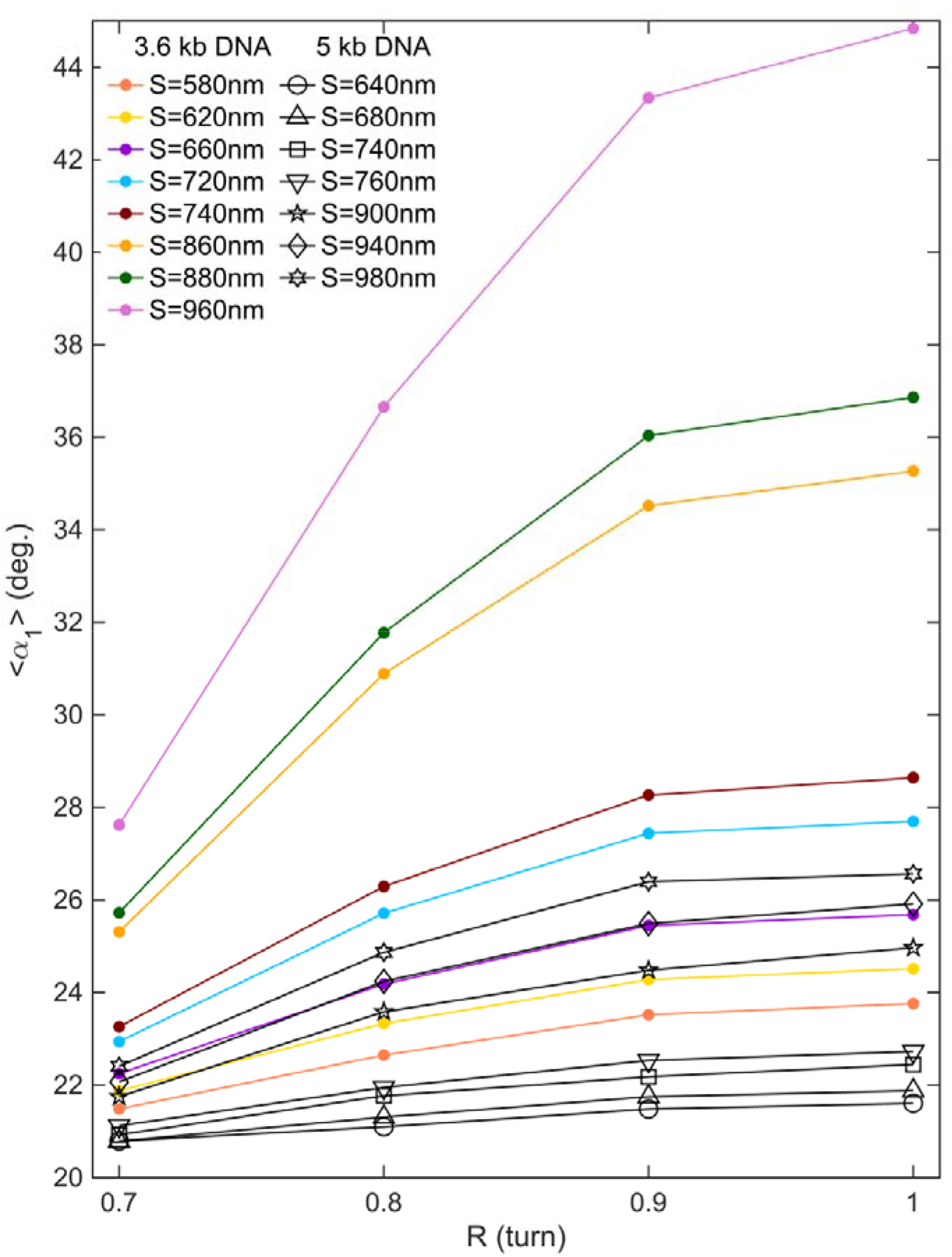
Comparison of crossing geometries achieved in single DNA-crossings for 3.6 kb and 5kb DNA. The mean values of bend angle, < _1_>, are plotted as a function of bead rotation (R) from simulations conducted at 2 pN force for different segment separation (S) values. The mean values from simulations performed at different segment separations are color coded for 3.6kb DNA (S = 580nm: Coral; 620 nm: Gold; 660 nm: Dark violet; 720 nm: Deep sky blue; 740 nm: Maroon; 860 nm: Orange; 880 nm: Dark green; 960 nm: Orchid) and are symbol coded for 5kb DNA (S = 640nm: Circle; 680 nm: Upward-pointing triangle; 740 nm: Square; 760 nm: Downward-pointing triangle; 900 nm: Pentagram; 940 nm: Diamond; 980 nm: Hexagram). The shorter 3.6 kb DNA molecules resulted in a much larger range of bend angles and “hookedness” as a function of bead rotation than the 5 kb DNA molecules used in previous unlinking measurements. The large variation in the imposed bend angles permits a sensitive measure of the relation between bend angle and topo IV unlinking rate.

### S11. Alternative analysis of simulations without parameterizing the crossings

The preferred crossings of topo IV obtained from the simulations were predicated on the geometric parameters defining the HJP model. To confirm our estimates of the preferred crossing geometry of topo IV, we performed an independent correlation analysis based on the pairwise distances between each bead in the pair of 5-bead arcs defining the crossing (Fig. S1). We considered the pairwise distance for 45 pairs in each crossing (Fig. S11). The correlation analysis of mean values of these pair-wise distances with unlinking rates gives a spectrum of correlation coefficients (Table S2). The strong anti-correlations (<-0.4) for pairs with non-central monomer beads are consistent with the weakly bent segments observed in the preferred crossing. Similarly, the moderate correlations for the intra-segment pair of neighboring beads to the central bead and the loss of correlation for the intra-segment pair of terminal beads do not support a sharp symmetric bend in the crossings. This approach offers an alternative, though qualitative, perspective on the preferred crossing geometry that is consistent with the preferred crossing geometry obtained through the correlation analysis.

**Figure S11:**
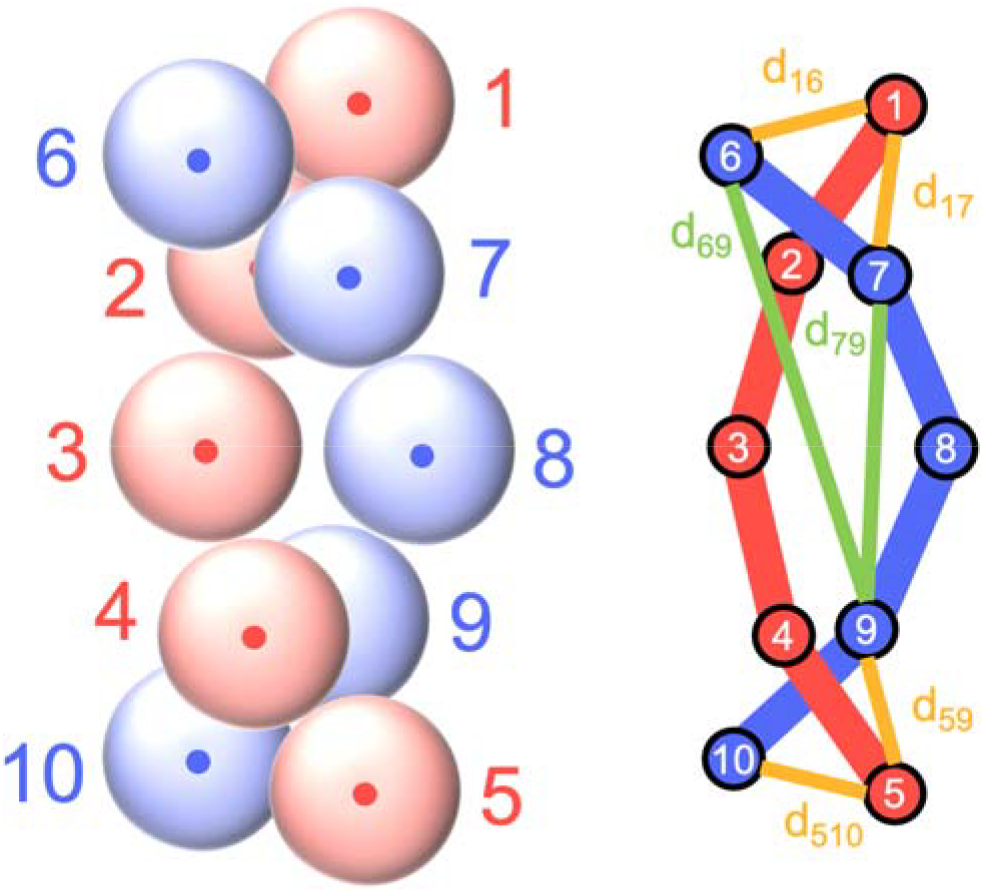
Schematic of a crossing of 5-monomer-bead segments from each dsDNA with the simplified ball-and-stick model (representing particle-and-bond) for easier visualization. The thin lines with labels show representative pair-wise distances.

### S12. Comparison of DNA bending energy for different bend angles

The bending energy of a DNA segment bent through a radius can be expressed as:

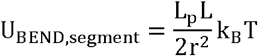

Where, L_p_ is the DNA persistence length, L is the DNA segment length, r is the radius of curvature of the segment, and k_B_T is the thermal energy. The bending energy can be expressed in terms of the bend angle (α) by replacing r with L/α in the above formula:

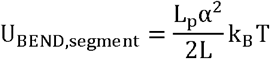

Hence, for a symmetric crossing with two uniformly bent segments,

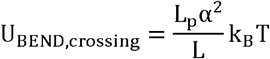

We consider L_p_ =50 nm, L=12.5 nm for further calculations

For the preferred crossing geometry (α_37°)

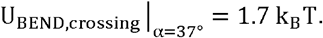

While, for the HJP crossing geometry (α _ 165°)

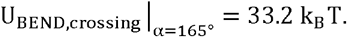

**Table S1:** Simulation parameters in the reduced Lennard–Jones (LJ) and SI units. The conversion of LJ units to SI units is not straightforward. Here, the LJ units are converted to SI units using the parameters provided on the LAMMPS website (https://docs.lammps.org/units.html).

|  | LJ units | SI units |
| --- | --- | --- |
| Length | $\sigma_{\text{LJ}}$ | 2.5 nm |
| Time | $\tau_{\text{LJ}}$ | 0.11 ns |
| Energy | $\epsilon_{\text{LJ}}$ | 4.14 pN nm |
| Force | $F_{\text{LJ}}$ | 0.61 pN |

**Table S2:** Correlation coefficients between unlinking rates and the mean values of different pair-wise distances in crossings. Column 1 some representative categories (color codes as Inter-segment pair: Orange; Intra-segment pair: Green) of the pairs shown in Columns 2 and 3 (color codes as Monomer from the first segment: Red; Monomer from the second segment: Blue).Columns 4 shows the Pearson correlation coefficient for corresponding unlinking rates. Columns 5 shows the associated p-values for the correlation calculations. The table shows results for 45 pairs in the ascending order of correlation coefficients.

| Pair type | ID_1 | ID_2 | PCC | p_value |
| --- | --- | --- | --- | --- |
| terminal | 1 | 6 | -0.50 | 0.0000 |
| terminal | 5 | 10 | -0.50 | 0.0000 |
|  | 1 | 7 | -0.45 | 0.0000 |
|  | 4 | 10 | -0.45 | 0.0000 |
|  | 2 | 6 | -0.45 | 0.0000 |
|  | 5 | 9 | -0.45 | 0.0000 |
|  | 2 | 7 | -0.42 | 0.0001 |
|  | 4 | 9 | -0.42 | 0.0001 |
| central | 3 | 8 | -0.37 | 0.0006 |
|  | 3 | 9 | -0.36 | 0.0007 |
|  | 2 | 8 | -0.36 | 0.0007 |
|  | 4 | 8 | -0.36 | 0.0007 |
|  | 3 | 7 | -0.36 | 0.0007 |
| central-terminal | 1 | 8 | -0.35 | 0.0010 |
| central-terminal | 3 | 10 | -0.35 | 0.0010 |
| central-terminal | 5 | 8 | -0.35 | 0.0010 |
| central-terminal | 3 | 6 | -0.35 | 0.0010 |
|  | 2 | 9 | -0.30 | 0.0064 |
|  | 4 | 7 | -0.30 | 0.0066 |
|  | 1 | 9 | -0.24 | 0.0286 |
|  | 2 | 10 | -0.24 | 0.0287 |
|  | 5 | 7 | -0.24 | 0.0292 |
|  | 4 | 6 | -0.24 | 0.0292 |
| trans terminal | 1 | 10 | -0.14 | 0.2078 |
| trans terminal | 5 | 6 | -0.14 | 0.2120 |
|  | 6 | 8 | -0.06 | 0.5793 |
|  | 8 | 10 | -0.06 | 0.5853 |
|  | 1 | 3 | -0.06 | 0.6007 |
|  | 3 | 5 | -0.06 | 0.6121 |
| terminal bend | 1 | 5 | 0.01 | 0.9095 |
| terminal bend | 6 | 10 | 0.01 | 0.9089 |
|  | 6 | 9 | 0.06 | 0.5979 |
|  | 2 | 5 | 0.06 | 0.5775 |
|  | 1 | 4 | 0.06 | 0.5701 |
|  | 7 | 10 | 0.06 | 0.5675 |
| bond | 4 | 5 | 0.08 | 0.4893 |
| bond | 6 | 7 | 0.08 | 0.4612 |
| bond | 8 | 9 | 0.10 | 0.3914 |
| bond | 1 | 2 | 0.10 | 0.3818 |
| bond | 9 | 10 | 0.11 | 0.3241 |
| bond | 7 | 8 | 0.12 | 0.2943 |
| bond | 2 | 3 | 0.13 | 0.2387 |
| bond | 3 | 4 | 0.14 | 0.2128 |
| mid bend | 7 | 9 | 0.28 | 0.0117 |
| mid bend | 2 | 4 | 0.29 | 0.0083 |

### Movie S1 (separate file)

Representative simulations movie illustrating crossings and its thermal motion at different degrees of hookedness. Low hookedness crossing (I): S = 440 nm, F = 0.5 pN, R = 0.7 turn (252°). Medium hookedness crossing (II): S = 860 nm, F = 1 pN, R = 0.9 turn (324°). High hookedness crossing (II): S = 960 nm, F = 2 pN, R = 1 turn (360°).

## Notes

https://doi.org/10.25444/nhlbi.28250900

### Competing Interest Statement

The authors have declared no competing interest.

